# The cost of fungicide resistance evolution in multi-field plant epidemics

**DOI:** 10.1101/2023.09.05.556392

**Authors:** Alexey Mikaberidze, Chaitanya S. Gokhale, Maria Bargués-Ribera, Prateek Verma

**Author notes:** Corresponding author 1. Corresponding author 2.

## Abstract

Epidemics of plant diseases are estimated to cause significant economic losses in crop production. Fungicide applications are widely used to control crop diseases but incur substantial indirect costs. One essential class of indirect costs arises due to the evolution of fungicide resistance. This indirect cost must be estimated reliably to design economic policy for more sustainable use of fungicides. Such estimation is difficult because the cost depends on economic parameters and the evo-epidemiological properties of crop pathogens. Even a conceptual framework for such estimation is missing. To address this problem, we combined a spatially implicit mathematical model of crop epidemics with an economic analysis at the landscape scale. We investigated how the net economic return from a landscape depends on the proportion of fungicide-treated fields. We discovered a pattern of accelerating (or decelerating) returns, contrary to expected diminishing returns. Next, we calculated the economic cost of the evolution of fungicide resistance as the difference between the optimal net return of the landscape in the absence and presence of resistance. We found that this cost depends strongly on the fungicide price, the degree of resistance, the pathogen’s basic reproduction number and the yield loss due to disease. Surprisingly, the cost declines with the fungicide price and exhibits a non-monotonic pattern as a function of the basic reproduction number. Hence, to calculate the cost, we must estimate these parameters robustly, incorporating variations in environmental conditions, crop varieties and the genetic composition of pathogen populations. Appropriate estimation of the cost of resistance evolution can inform economic policy and encourage more sustainable use of fungicides.

**Author Summary:** Fungicides protect crops from diseases and are essential for securing the global food supply, but they incur serious indirect costs to society and the environment. One such cost arises because of the evolution of fungicide resistance. Part of the pathogen population (fungicide-resistant) can gain protection from a fungicide via a genetic change. If fungicide applications continue, the fungicide-resistant subpopulation increases and dominates the population, leading to low efficacy. Resistance can lead to severe economic losses, but no conceptual framework exists for estimating them. We present a novel mathematical framework to estimate the economic costs of fungicide resistance at the landscape scale. We combined an epidemiological model describing disease spread with an economic cost-benefit analysis. Surprisingly, we found that the economic cost of resistance declines for more expensive fungicides. This cost also depends on the pathogen’s capacity to spread (invasiveness): the cost is highest for pathogens with intermediate invasiveness. Thus, the cost of resistance depends on economic parameters and the biological characteristics of plant diseases. Our findings can inform economic policies for sustainable fungicide use, such as taxes or subsidies. Our paper also contributes to the broader discourse on agricultural sustainability while ensuring global food security.

## 1 Introduction

Epidemics of plant diseases are estimated to cause significant economic losses in crop production (Ostfeld et al., 2005; Oerke, 2006; Savary et al., 2019; Global Plant Health Assessment, 2023). To reduce losses, farmers frequently apply fungicides. They can maximize the net economic return of fungicide applications, e.g., by optimizing the fungicide dose based on the balance between the yield benefit and the cost of fungicide application (Jørgensen et al., 2017; Hillebrandt, 1960). However, this approach is problematic for at least two reasons.

First, fungicide applications not only have direct economic costs, but they also incur considerable indirect costs, such as environmental costs, human health costs, and costs associated with the evolution of fungicide resistance (Pimentel et al., 1980; Pimentel, 2005; Bourguet and Guillemaud, 2016). One opportunity to manage crop diseases in a more sustainable manner is to incorporate indirect costs, at least partially, into fungicide prices. This opportunity can be realized using economic policy instruments, e.g., by introducing special pesticide taxes or subsidies, as has been done in several countries, including France, Denmark, Norway and Sweden (Finger et al., 2017; Lee et al., 2019). However, it is not easy to design pesticide tax rates or subsidies to reflect indirect costs, as indirect costs are notoriously difficult to estimate (Pimentel et al., 1980; Pimentel, 2005; Bourguet and Guillemaud, 2016).

Second, many crop pathogens of economic relevance can disperse over long distances, not only within individual fields but also between fields and across entire regions (Brown and Hovmoller, 2002). Therefore, decisions regarding disease management made at a particular farm can affect epidemic development in other farms across a region, and decisions that are optimal for an individual farm may turn out to be sub-optimal at the scale of a regional cultivated landscape. This is also the case when managing weeds: for instance, Evans et al. (2018) used computational modeling to show that aggregating the best herbicide resistance management practices at the landscape scale can slow down the evolution of resistance.

An essential class of indirect costs of fungicide applications arises because pathogen populations can adapt to fungicides via the evolution of fungicide resistance, which reduces fungicide efficacy (Fisher et al., 2018). From an economic perspective, these costs represent an externality (Ayres and Kneese, 1969): development of resistance is caused by fungicide applications on specific fields, but resistance can spread between fields and across farms and entire regions. Hence, a much wider community of farmers will eventually suffer the economic costs of fungicide resistance evolution.

These costs are difficult to estimate because they depend on the economic parameters of the affected cropping system, the epidemiological, and the evolutionary properties of a particular pathosystem. A phone survey of 137 growers in Western Australia’s Wheatbelt revealed that growers spent, on average, AU$42 per hectare on fungicide treatments of barley diseases in the 2019/2020 growing season. The growers were further willing to spend on average AU$18 per hectare to delay or mitigate fungicide resistance (Olita et al., 2023). The latter estimate represents a perceived economic cost of fungicide resistance, and it remains to be investigated to what extent this perception corresponds to the actual cost. At a wider national scale in the USA, the economic impact of pesticide resistance was estimated at US$1.5 billion in 2005, which, when adjusted for inflation, amounts to U$2.5 billion today, excluding the rising costs associated with increased pesticide use (Beckerman et al., 2023). A national-scale investigation of herbicide resistance in black-grass (*Alopecurus myosuroides*) across the UK indicated that herbicide resistance can double the economic costs of weed management (Hicks et al., 2018). Rough estimates of economic losses in crop production due to insecticide resistance were in the range 10-15 % (Pimentel et al., 1980; Pimentel, 2005). These were based on economic data on cotton production in the USA, assuming that insecticide use increases due to resistance and that this increase is the only cost incurred by resistance. However, it is unclear to what extent this assumption is fulfilled; it is also difficult to extrapolate these estimates to other crops and regions and extrapolate from insecticide resistance to fungicide resistance. Only a few attempts have been made to estimate the economic costs of fungicide resistance in crop pathogens in the literature, and these have focused on individual fields (van den Bosch et al., 2018, 2020). Even a conceptual framework to enable the calculation of economic costs of fungicide resistance across wider spatial scales is lacking. To fill this knowledge gap, we addressed the following questions: (i) “How to calculate the economic cost of fungicide resistance?” and (ii) “How does the economic cost of fungicide resistance depend on the economic and epidemiological/evolutionary parameters of the crop-pathogen system?”

For this purpose, we combined a spatially implicit epidemiological model of crop epidemics with an economic analysis at the landscape scale (bioeconomic modeling). Using this approach, we calculated the economic cost of the evolution of fungicide resistance, *C*_*R*_, as the difference between the optimal net return of the landscape in the absence and presence of resistance. This allowed us to explore how *C*_*R*_ depends on the key epidemiological/evolutionary and economic parameters of the system. Our results provide a conceptual basis for the estimation of *C*_*R*_ as a key component of indirect costs of fungicide applications, which can inform economic policy to achieve a more sustainable use of fungicides.

## 2 Materials and methods

### 2.1 Epidemiological model for multiple fields

We study the epidemiological dynamics of a generic fungal pathogen of crop plants at the scale of multiple fields (Fig. 1). We consider a regional cultivated landscape composed of *N* fields growing the same annual crop (e.g., wheat or maize). The key variables and parameters are given in Table 1. First, we devise a model considering a wildtype (fungicide-sensitive) pathogen strain with a fungicide treatment. Then, we extend this model to incorporate a fungicide-resistant pathogen strain.

**Table 1:** Key variables and parameters of the model.

| Symbol | Description |
| --- | --- |
| **Variables** |  |
| $H_i$ | Number of healthy fields with treatment status $i$ ($i = u$ for untreated, $i = t$ for treated) |
| $I_{ij}$ | Number of fields infected by pathogen strain $j$ ($j = w$ for wildtype, $j = r$ for resistant) with treatment status $i$ ($i = u$ for untreated, $i = t$ for treated) |
| **Parameters** |  |
| $N$ | Total number of fields |
| $\theta$ | Fraction of fungicide-treated fields (or fungicide coverage); 0-1 |
| $\epsilon_i$ | The efficacy of fungicide treatment in reducing the transmission rate of the pathogen strain $i$ ($i = w$ for wildtype, $i = r$ for resistant); 0-1 |
| $\beta_i$ | Transmission rate of pathogen strain $i$ ($i = w$ for wildtype, $i = r$ for resistant) from infected to the healthy field |
| $\mu$ | Recovery rate of infected fields to become healthy fields |
| $c$ | The price of fungicide application |
| $y_H$ | Yield from a healthy field per season |
| $f$ | Relative fungicide price with respect to yield from a healthy field |
| $y$ | Relative yield of a diseased field with respect to the yield from a healthy field |
| $Y$ | Total yield of the landscape (expressed as a fraction of the monetary equivalent of the maximum yield) |
| $g(\theta)$ | Net return relative to yield from a healthy field, $y_H$ |
| $\theta^*$ | Optimal fraction of treated fields that maximizes $g(\theta)$ |

**Figure 1:**
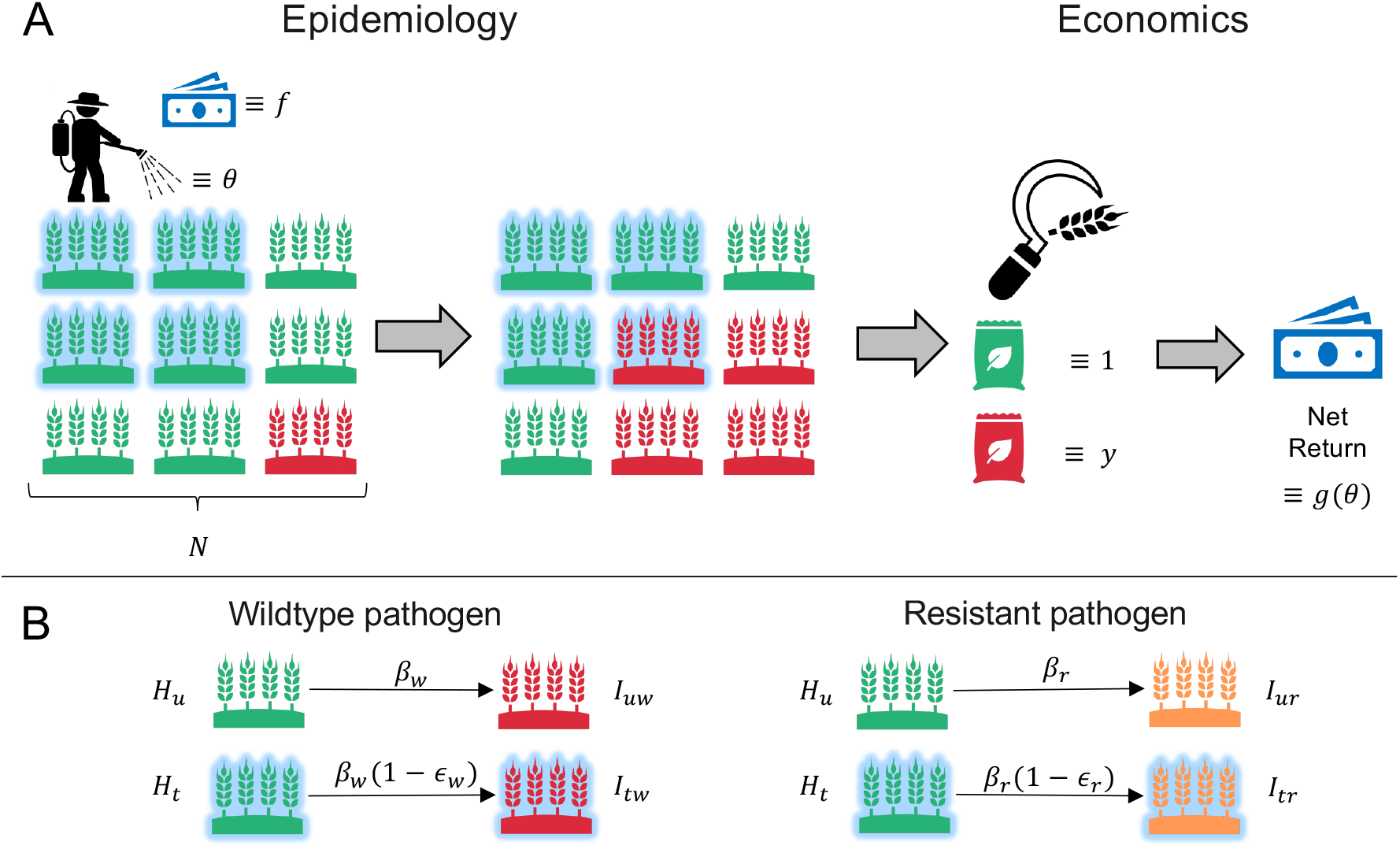
Overview of the multiple field model of plant disease incorporating both epidemiology and economics aspects in the presence of fungicide treatment. Healthy fields (*H*_*u*_ and *H*_*t*_) are shown in green, fields infected with the wildtype pathogen strain (*I*_*uw*_ and *I*_*tw*_) are shown in red, fields infected with the resistant pathogen strain (*I*_*ur*_ and *I*_*tr*_) are shown in orange. Fungicide-treated fields have a light-blue glow around them. (A) Out of the total *N* fields, initially, only one is infected, while the rest are healthy. A fraction *θ* of all fields are treated with a fungicide, which comes at a price *f*. The net return (*g*(*θ*)) of the harvest is calculated as the monetary equivalent of the yield from all healthy and infected fields (1 and *y*, respectively) minus the total fungicide price. (B) The state variables of the epidemiological model and the transitions between them: healthy fields can become infected over time.

#### 2.1.1 Model with fungicide treatment

Each field can be either healthy (*H*) or infected (*I*) with a wildtype pathogen (subscript *w*), and each field can be treated (subscript *t*) or untreated (subscript *u*) with the fungicide. Hence, we have four possible field states: *H*_*u*_, *H*_*t*_, *I*_*uw*_, and *I*_*tw*_. *H*_*u*_ and *I*_*uw*_ represent the numbers of untreated healthy and infected fields, respectively, while *H*_*t*_ and *I*_*tw*_ represent the treated fields. The dynamics of the disease are captured by equations that are similar to the epidemiological susceptible-infected model:

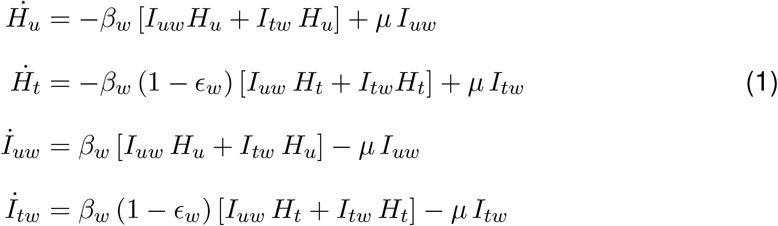

The wildtype pathogen strain spreads from infected fields to healthy fields at a rate *β*_*w*_. In treated fields, this rate is reduced by a factor 1 − *ϵ*_*w*_, where *ϵ*_*w*_ is the fungicide efficacy. Infected fields recover and convert back to healthy at a rate *µ*. The fraction *θ* = (*H*_*t*_ + *I*_*tw*_)*/N* of all fields is treated with fungicide. Hence, the number of untreated and treated fields can be calculated as,

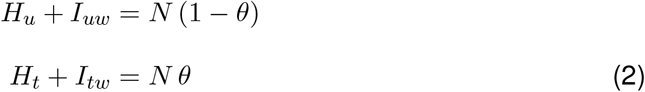

First, consider the no-treatment scenario, *θ* = 0, for the calculation of the basic reproduction number. It can be interpreted as the average number of secondarily infected fields produced by a single infected field introduced into a population of healthy fields. The basic reproduction number (*R*_0_) for no-treatment scenario is,

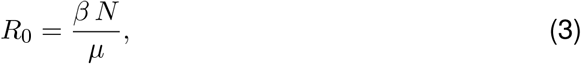

as derived in Subsec. A.1 in S1 Appendix. A higher value of *R*_0_ indicates a higher potential for the pathogen to spread.

When we include fungicide treatment (*θ >* 0), we use the next-generation matrix method in Subsec. A.2.3 in S1 Appendix (Diekmann et al., 1990; van den Driessche, 2017) to derive the effective reproduction number as

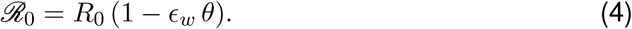

This expression quantifies how fungicide treatment affects the pathogen’s capacity to invade and spread. The critical fungicide coverage, for which *R*_0_ = 1 can be calculated as:

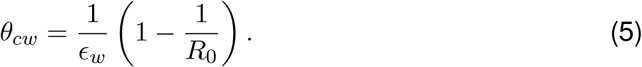

Treating the fields beyond the critical coverage level (*θ > θ*_*cw*_) prevents an epidemic. This criterion depends on the fungicide efficacy, *ϵ*_*w*_, and the pathogen’s *R*_0_ without treatment. Thus for a given *R*_0_, when *θ > θ*_*cw*_, fungicide treatment will suppress *R*_0_ to a value below one if *ϵ*_*w*_ is high enough, i.e. if *ϵ*_*w*_ *> ϵ*_*cw*_ = 1 − 1*/R*_0_. However, at a low fungicide efficacy *ϵ*_*w*_ *< ϵ*_*cw*_, even treating all fields (*θ* = 1) will not prevent an epidemic as *R*_0_ will remain higher than one. Similarly, for a fungicide with a given efficacy *ϵ*_*w*_, increasing *θ* to values above the critical, *θ*_*cw*_, will drive the pathogen to extinction only for pathogens with *R*_0_ *< R*_0*cw*_, where

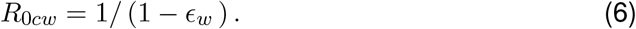

The expressions for the critical *R*_0*cw*_ and *θ*_*cw*_ will help us to better understand and perform bioeconomic analysis below in Sec. Bioeconomic analysis of fungicide use. However, to conduct a bioeconomic analysis, we must consider the evolutionary aspect of the problem as repeated fungicide application can lead to the evolution of resistance in pathogen populations (Mikaberidze et al., 2014, 2017; Fisher et al., 2018), which we explore next.

#### 2.1.2 Model with fungicide treatment and resistance

We extend the model in Eqs. (1) to incorporate a resistant pathogen strain that could be fully or partially protected from the fungicide. We assume that either a wildtype (*w*) or a resistant (*r*) pathogen strain dominates each infected field (Parnell et al., 2006). The epidemiological dynamics are given by:

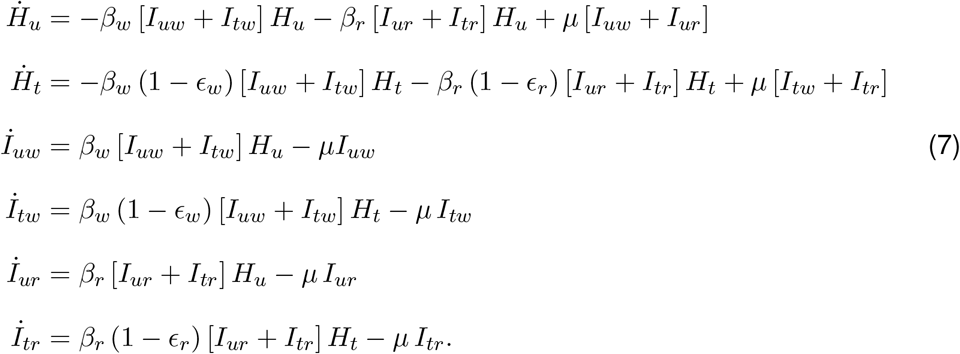

Resistance reduces fungicide efficacy. Consequently, fungicide efficacy against the resistant strain is lower than its efficacy against the wildtype strain, *ϵ*_*r*_ *< ϵ*_*w*_. Further, the transmission rate of the resistant strain, *β*_*r*_, can be reduced because of possible fitness costs associated with resistance mutations. We neglect this effect here (i.e., set *β*_*r*_ = *β*_*w*_) but consider it in Subsec. B.2.2 in S1 Appendix. The fraction of treated fields is given by *θ* = (*H*_*t*_ + *I*_*tw*_ + *I*_*tr*_)*/N*. The stability analysis for the Eqs. (7) is presented in Subsec. A.3.1 - A.3.3 in S1 Appendix.

Using the next-generation matrix approach, we derive the following expression for the effective reproduction number in the more general case of a partially effective fungicide and partial resistance (Subsec. A.3.4 in S1 Appendix):

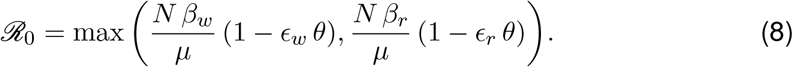

We consider equal transmission rates for the wildtype and resistant strains (*β*_*w*_ = *β*_*r*_) but unequal fungicide efficacies for the two strains (*ϵ*_*w*_ *> ϵ*_*r*_). As a result, the second expression on the right-hand side of Eq. (8) is larger than the first expression, and the effective reproduction number takes the form:

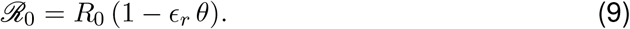

The above expression is similar to Eq. (4) but has *ϵ*_*r*_ instead of *ϵ*_*w*_ as a factor reducing the basic reproduction number. Based on Eq. (9), we determine the critical fungicide coverage, above which the fungicide treatment leads to the extinction of resistance:

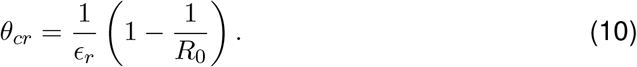

If *θ*_*cr*_ *>* 1, then the fungicide treatment cannot drive the pathogen population (consisting of the resistant strain) to extinction even when fungicide coverage is complete *θ* = 1. This is the case when the fungicide efficacy against the resistant strain is lower than the critical value, *ϵ*_*r*_ *< ϵ*_*cr*_, where

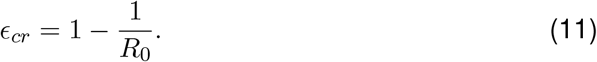

For a fungicide with a given efficacy *ϵ*_*r*_ against the resistant strain, increasing the fraction of treated fields *θ* to values above the critical, *θ*_*cr*_, can drive the pathogen to extinction for a sufficiently low basic reproduction number, *R*_0_ *< R*_0*cr*_, where

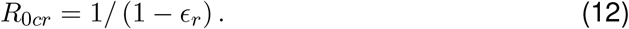

### 2.2 Methodological features and assumptions

Our model assumes density-dependent transmission or pseudo-mass-action dynamics (Eq. (7)) (Keeling and Rohani, 2008). Disease transmission increases with the density of fields, and there is no explicit spatial structure. The approximation works well when the primary mode of long-distance dispersal of plant pathogens is via networks of transportation/trade (e.g., via movement of infected plants or other materials). For plant pathogens that disperse via air-borne spores, a spatially explicit model would need to consider a distance-dependent transmission (Medlock and Kot, 2003; Mikaberidze et al., 2016) that has been characterized empirically in a number of important crop diseases (e.g., by Karisto et al. (2022, 2023)). Our model provides a valuable reference scenario, but is expected to overestimate the number of diseased fields at endemic equilibrium.

Although our model considers multiple growing seasons, we simplified it as in Parnell et al. (2006) by neglecting the cyclic nature of multi-seasonal dynamics due to, e.g. the periodic absence of host plants. Environmental stochasticity associated with such multi-seasonal dynamics may play a role in the emergence of novel, better-adapted pathogen genotypes (Willocquet et al., 2020).

The model belongs to the class of spatially implicit metapopulation models and is similar to the model developed previously (Parnell et al., 2006). Similarly to Parnell et al. (2006), we assume that environmental conditions and cropping practices are the same in different fields and that all fields have the same size. Hence, the model parameters do not vary between different fields. We set possible states for each field to be either healthy or infected (and infectious). Although in reality, the level of infection is continuous, our assumption implies that each infected field produces a similar amount of inoculum that can be transmitted to other fields. Thus, the characteristic time scale of within-field epidemic development is much shorter than the time scale of regional-scale dynamics. This idealized scenario is likely to be closer to reality for foliar fungal diseases of field crops caused by pathogens with long-distance dispersal in regional cultivated landscapes with relatively uniform environments and where crop cultivars have a similar degree of susceptibility to the disease. Examples of such diseases and cropping systems include northern corn leaf blight disease of maize in the corn belt regions of the US; target spot in soybean in major soybean-producing areas of Brazil (e.g., Mato Grosso); and wheat rusts (e.g., stripe rust and leaf rust) in major wheat-producing areas of the US (e.g., Central Great Plains). It will be interesting to relax this assumption in future work, allowing for heterogeneous disease levels among different fields, and investigate how this heterogeneity affects the conclusions of this study.

The model assumes that recovered fields are susceptible to re-infection (i.e., no induced resistance) and neglects demographic stochasticity (Mikaberidze et al., 2017). However, in contrast to Parnell et al. (2006), our model does not have an explicit spore compartment, assuming that dispersal occurs much faster than other relevant epidemiological processes. We also assume that a field can be infected by either a sensitive or a resistant pathogen strain, but not by both. We justify this assumption similarly to Parnell et al. (2006). When a resistant strain appears in a fungicide-treated field, it can invade and outcompete the sensitive strain if the selection is strong enough and the fitness cost of resistance is low enough (Mikaberidze et al., 2014). This condition is often fulfilled, as the fitness costs are typically below 10% (Mikaberidze et al., 2015; Hawkins and Fraaije, 2018), and hence most resistant strains, even those carrying partial resistance, will have a substantial fitness advantage over the sensitive strains in fungicide-treated fields. Alternatively, the resistance strain can fail to invade and die out or remain at a low level according to mutation-selection or migration-selection balance. In the latter scenario, the resistant strain does not affect the fungicide efficacy and is unlikely to spread to other fields, and therefore, the assumption that the field is infected only by the sensitive strain is justified.

The above assumptions (along with their caveats) allowed us to formulate a simple landscape-scale model of a generic fungal disease of crop plants. We can then obtain analytical outcomes and capture essential features of disease dynamics across entire ranges of plausible parameter values. To describe specific crop-pathogen systems with a higher degree of biological realism, our model will need to relax some of the assumptions at the expense of increased model complexity.

Having incorporated fungicide resistance into the model, we now analyze the economic aspects of fungicide treatment. We aim to determine the optimal fungicide treatment coverage that maximizes the net economic return.

### 2.3 Economic analysis: Net return and optimal fungicide coverage

We define the net return as the total income resulting from the sales of the harvested yield minus the cost of fungicide treatment across all *N* fields in the landscape over the course of *K* growing seasons. Mathematically, the net return is:

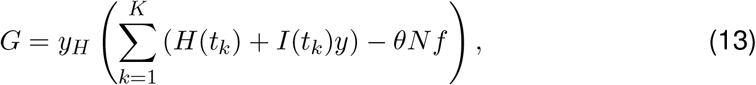

where *y*_*H*_ is the yield of a healthy field, *t*_*k*_ is the time point corresponding to the end of season *k, y* = *y*_*I*_*/y*_*H*_ is the relative yield of a diseased field (hence, the relative yield loss is 1 − *y*), *f* = *c/y*_*H*_ is the relative fungicide price. Here, *c*, the cost of fungicide application per field, includes the cost of spraying material and application cost. The total yield of the landscape, *Y*, is given by the sum of the first and the second terms in Eq. (13): 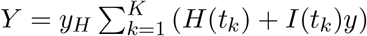; and the total fungicide price over the landscape, *F*, is given by the third term in Eq. (13): *F* = *y*_*H*_*θNf*.

The net return does not change after the system reaches a stable equilibrium. Besides being analytically tractable, we consider this case as it corresponds to the limit of sustainable disease management. The outcomes will likely remain the same across a range of times, even before the equilibrium is strictly reached. However, they are expected to be different during the initial stages of dynamics as these are affected by initial conditions.

At equilibrium, we characterize the net return as follows. We compute it for a single season and divide it by the total number of fields *N* to obtain the average across fields. We also divide it by yield per season from a healthy field, *y*_*H*_, to obtain the relative net return (which we will call “net return” for brevity) *g*(*θ*) = (*H*^∗^ + *I*^∗^*y*) */N* −*θf*, where 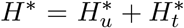 and 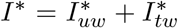 represent the values of the state variables (numbers of healthy and diseased fields) at a stable fixed point. The yield from healthy and diseased fields boosts the net return (the first term), while the cost of fungicide treatment reduces the net return (the second term). Since the total number of fields remains constant, *H*^∗^ = *N* − *I*^∗^, we rewrite the expression for the net return in the form that supports further analyses:

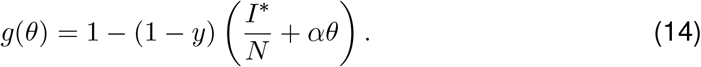

Here, the net return without disease and without treatment is equal to one (the first term in Eq. (14)). The second term in Eq. (14) quantifies the reduction of the net return due to disease, which is proportional to the fraction of diseased fields *I*^∗^*/N*. The third term in Eq. (14) quantifies the reduction of the net return due to the cost of fungicide treatment. This term is proportional to *θ* with the coefficient *α* that we call the cost ratio parameter:

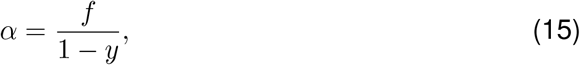

where *f* is the relative fungicide price, and 1 − *y* is the yield loss in a diseased field (expressed as its monetary equivalent). The proportion of diseased fields 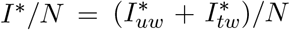 in the second term also depends on *θ* albeit in a complex manner according to Eqs. (A4-A11) (Subsec. A.2 in S1 Appendix) and the associated linear stability conditions.

We define the optimal fungicide coverage or simply optimal coverage, *θ*^∗^, as the proportion of treated fields that results in the highest net return. The optimal coverage *θ*^∗^ can be determined by calculating the value of *g*(*θ*) across a fine grid of *θ*-values.

The *θ*-value that maximizes *g*(*θ*) is the optimal coverage *θ*^∗^. In cases where multiple values of *θ* yield the same highest value of *g*(*θ*), we define the lowest among these as the optimal coverage *θ*^∗^.

### 2.4 Net return in an individual field

While the focus of this work is on cultivated landscapes with multiple fields, we provide the model for the net return in an individual field for comparison. When managing disease in an individual field, growers can adjust the fungicide dose to maximize the net return, *g*, given by

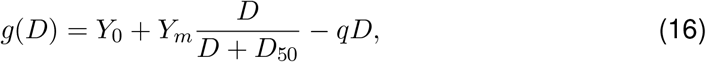

where *Y*_0_ is the yield without fungicide application, and *D* is the fungicide dose. The second term in Eq. (16) is the yield benefit of the fungicide application, where the functional form used is a simplification of the Hill function, which describes a typical empirical dose-response relationship for fungal diseases in cereal crops (Mikaberidze et al., 2017). Hence, the total yield of a field, *Y*, is given by the sum of the first and second terms in Eq. (16): 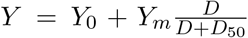. (We also defined *Y* above as the total yield of a landscape; to avoid confusion, we specify the context every time we use *Y*). The parameter *Y*_*m*_ represents the maximum effect at large doses, and *D*_50_ corresponds to the dose for which half of the maximum effect is achieved. The third term in Eq. (16) quantifies the total fungicide price, *F* = *qD*, where *q* is the price per unit fungicide. Net return, *g*(*D*), in Eq. (16) exhibits a maximum at the fungicide dose

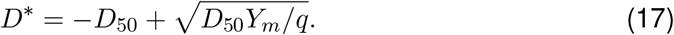

If the fungicide becomes sufficiently expensive (*q > Y*_*m*_*/D*_50_), the expression for the optimum dose in Eq. (17) becomes negative, meaning that the net return is maximized when no fungicide is applied. For less expensive fungicides, the net return does exhibit a maximum at positive doses. The optimum dose is higher for cheaper fungicides. We refer the reader to Subsec. B.1 in S1 Appendix for more details on the individual field model.

## 3 Results

### 3.1 Bioeconomic analysis of fungicide use

To perform a bioeconomic analysis, we first set the context by comparing the optimal fungicide treatments between individual fields and cultivated landscapes. Next, we answer the central question: What fraction of fields should be treated with a fungicide to maximize the net return of a landscape?

#### Optimal fungicide treatments in individual fields versus cultivated landscapes

When controlling disease in an individual field, the yield, *Y* (expressed as its monetary equivalent), typically exhibits a saturating increase as we increase the fungicide dose, *D* (second term in Eq. (16); Fig. 2A). The total fungicide price, *F*, increases linearly when increasing the fungicide dose with the slope given by the fungicide price per unit dose (third term in Eq. (16); dark red lines in Fig. 2A). The net return, *g*(*D*), is then given by the difference between the yield, *Y*, and the total fungicide price, *F* (Eq. (16), Fig. 2A). *g*(*D*) reaches a maximum for an intermediate (optimal) fungicide dose. For doses higher than optimal, the cost associated with increasing the dose is no longer compensated by an associated increase in yield. These relationships follow “the law of diminishing returns” that has achieved an almost universal status in the economics of crop production (Kho, 2000) and other areas of economics (Shephard, 1970; Brue, 1993).

**Figure 2:**
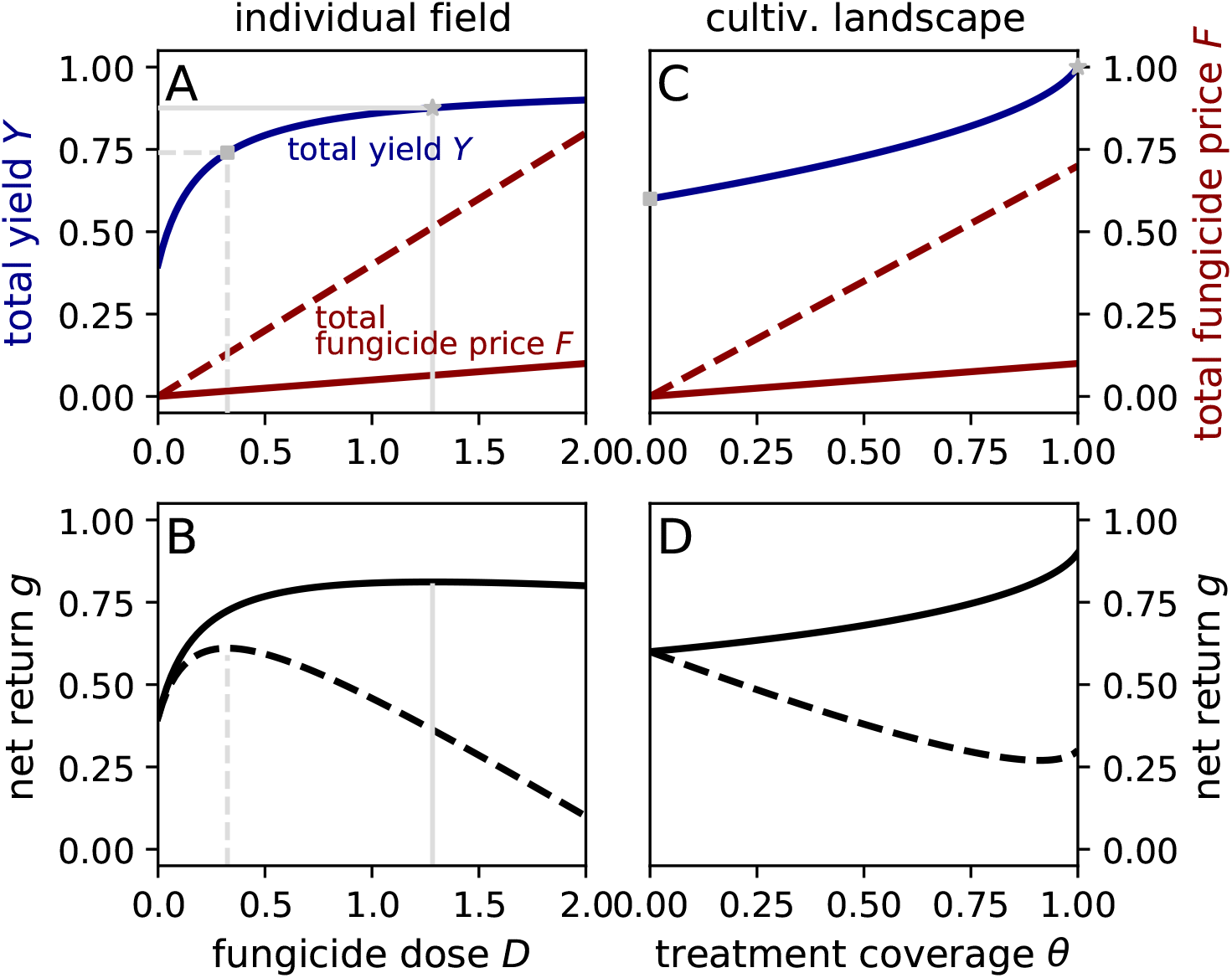
How does the net return depend on the intensity of treatment? (A) In an individual field, the yield *Y* exhibits a saturating increase as we increase the fungicide dose, *D* [solid blue; Eq. (B2) in S1 Appendix]. *Y* is expressed as a fraction of the monetary equivalent of the maximum yield. The total fungicide price, *F*, is proportional to the dose, *D*, with the slope given by the price per unit: *q* = 0.05 (solid red); *q* = 0.4 (dashed red). (B) Black curves represent the net return computed as the blue curve in (A) minus one of the red lines in (A), with *q* = 0.05 (solid) and *q* = 0.4 (dashed). The net return exhibits a maximum for an intermediate dose [Eq. (B4) in S1 Appendix]. Grey vertical lines in (A) and (B) show doses that maximize the net return. Other parameter values for (A) and (B): *y*_*m*_ = 0.55, *D*_50_ = 0.2, *y*_0_ = 0.4. (C) In a cultivated landscape, the total yield of the landscape, *Y*, [Eq. (B8) in S1 Appendix] exhibits an accelerating increase as we increase the fraction of treated fields, *θ* (blue). The total fungicide price, *F*, is proportional to *θ*, with the slope given by the relative fungicide price: *f* = 0.1 (solid red), *f* = 0.7 (dashed red). (D) As a result, the net return is maximum when treating either all fields (*θ* = 1, solid, *f* = 0.1) or when not treating any fields (*θ* = 0, dashed, *f* = 0.7); according to Eq. (B7) in S1 Appendix. Other parameter values for (C) and (D): *β* = 0.025, *µ* = 5, *N* = 1000 (hence, *R*_0_ = *βN/µ* = 5), *y* = 0.5.

In contrast, when controlling disease in a cultivated landscape, the total yield of the landscape (*Y*, expressed as its monetary equivalent, Eq. (B8) in S1 Appendix exhibits an accelerating increase as we increase the fungicide coverage, *θ* (Fig. 2C). The total price of the fungicide treatment increases linearly with *θ* (Fig. 2C), as it did for an individual field above. The net return, *g*(*θ*), is again the difference between the monetary yield, *Y*, and the total fungicide price, *F*. However, here, the outcome is very different. The net return does not exhibit a maximum at any intermediate *θ* value. Thus, the net return can be maximized only at the extreme values of *θ*: *θ* = 0 treating no fields or *θ* = 1 treating all fields. When the fungicide is sufficiently cheap, *θ* = 1 maximizes the net return (Fig. 2C,D). However, when the fungicide is expensive, then the no-treatment strategy (*θ* = 0) maximizes the net return (Fig. 2C,D).

Thus, the law of diminishing returns does not necessarily hold when controlling disease in cultivated landscapes. This finding has important implications for the economics of crop disease management, optimal decision making and associated policy-making, which is often done at scale and not for individual fields.

#### Optimal fungicide coverage without resistance

Our analysis reveals that three parameters influence the optimal fungicide coverage, *θ*^∗^: (i) the fungicide price, *f*, (ii) the yield loss in a diseased field, 1 − *y* and (iii) the critical fraction of treated fields, *θ*_*cw*_ (in Eq. (5)). The first two parameters affect *θ*^∗^ only through their ratio, the cost ratio parameter *α* = *f/*(1 − *y*) (defined in Eq. (15) above). Therefore, below, we study the effects of *α* and *θ*_*cw*_.

When *α* is sufficiently low, treating all fields (*θ*^∗^ = 1, if *R*_0_ *> R*_0*cw*_; Fig. 3A) or most fields (*θ*^∗^ = *θ*_*cw*_, if *R*_0_ *< R*_0*cw*_ Fig. 3A) maximizes the net return (here, *θ*_*cw*_ is given by Eq. (5)). When *α* is sufficiently high, not treating any fields maximizes the net return (*θ*^∗^ = 0; Fig. 3A,B). Thus, the net return can only be maximized at extreme values of the fungicide coverage, either *θ*^∗^ = 0 or *θ*^∗^ = 1 (or *θ*^∗^ = *θ*_*cw*_ in case *θ*_*cw*_ *<* 1, whereby *θ*_*cw*_ is still close to 1).

**Figure 3:**
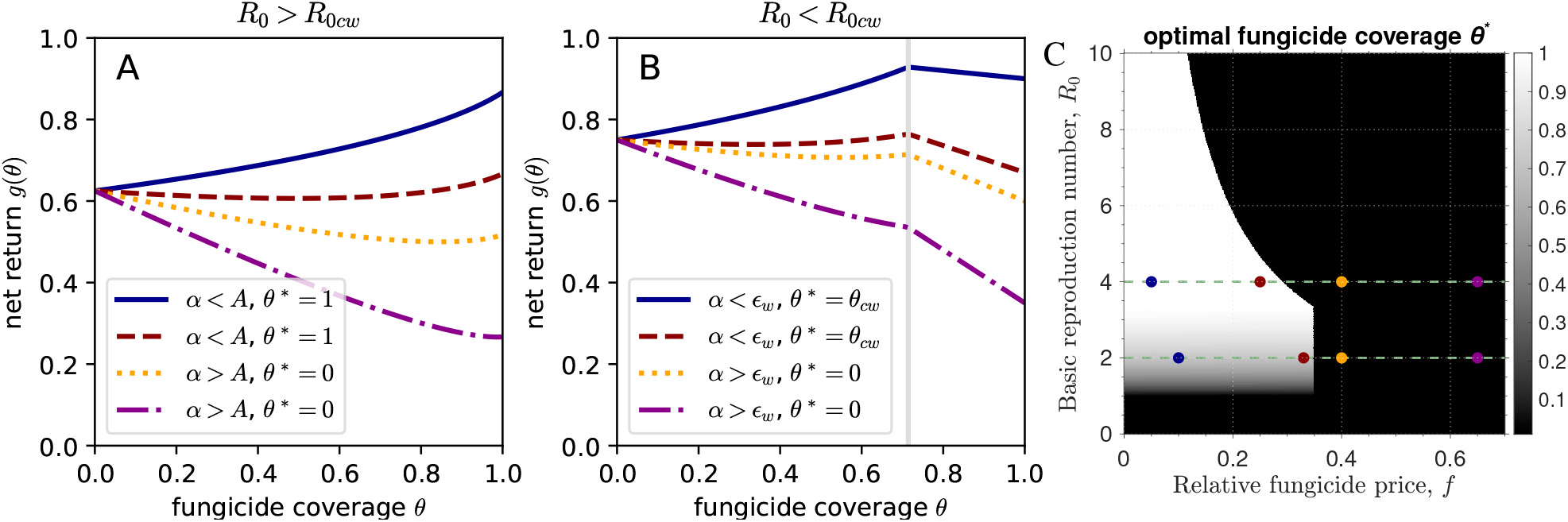
What is the optimal fungicide coverage in the landscape model without fungicide resistance? (A) The net return *g*(*θ*) is plotted against the fungicide coverage (fraction of treated fields *θ*) for *R*_0_ *> R*_0*cw*_ and *θ*_*cw*_ *>* 1 (i.e., *β* = 0.02 and *R*_0_ = *βN/µ* = 4). The four curves represent different relative fungicide prices (*f*): 0.05 (solid blue), 0.25 (dashed red), 0.4 (dotted orange), and 0.65 (dash-dotted magenta). (B) The net return *g*(*θ*) is plotted against *θ* for *R*_0_ *< R*_0*cw*_ and *θ*_*cw*_ = 0.714 *<* 1 (shown by a vertical grey line, i.e., *β* = 0.01 and *R*_0_ = *βN/µ* = 2). The four curves represent different values of *f* : 0.1 (solid blue), 0.33 (dashed red), 0.4 (dotted orange), and 0.65 (dash-dotted magenta). (C) The heatmap shows the optimal fungicide coverage *θ*^∗^ for varying fungicide price *f* and basic reproduction number *R*_0_. In the white region, treating all fields is optimal; in the black region, not treating any fields is optimal; in the grey region, treating only a fraction of fields is optimal. Each curve in (A) and (B) corresponds to a point on the green dashed lines in the heatmap (C). These points are highlighted by circles of the same color as lines in (A) and (B). Specifically, *R*_0_ = 4 and *R*_0_ = 2 correspond to (A) and (B) respectively. Other parameter values are constant for all panels: *N* = 1000, *µ* = 5, *ϵ*_*w*_ = 0.7, and *y* = 0.5.

To gain a quantitative understanding of factors affecting the optimal coverage, *θ*^∗^, consider the regimes *θ*_*cw*_ *>* 1 (*R*_0_ *> R*_0*cw*_) and *θ*_*cw*_ *<* 1 (*R*_0_ *< R*_0*cw*_) in more detail. When *θ*_*cw*_ *>* 1, it is impossible to wipe out the epidemic even if we treat all fields. This is the case for pathogens with a sufficiently high *R*_0_, whereby, *R*_0_ *> R*_0*cw*_ (Fig. 3A), where *R*_0*cw*_ is given by Eq. (6). On the contrary, when *θ*_*cw*_ *<* 1, one can wipe out the epidemic by treating a high enough proportion of fields, i.e., *θ > θ*_*cw*_. This is the case for pathogens with a sufficiently low *R*_0_, whereby, *R*_0_ *< R*_0*cw*_ (Fig. 3B).

For *θ*_*cw*_ *>* 1 (*R*_0_ *> R*_0*cw*_), not treating any fields (*θ* = 0) maximizes the net return when the cost ratio parameter is sufficiently high (*α > A*). In contrast, when *α < A* treating all fields (*θ* = 1) maximizes the net return (Fig. 3A), where

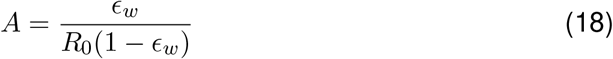

is a combination of epidemiological parameters.

For *θ*_*cw*_ *<* 1 (*R*_0_ *< R*_0*cw*_), the optimal coverage, *θ*^∗^, depends on how the cost-ratio *α* relates to the fungicide efficacy *ϵ*_*w*_. If *α > ϵ*_*w*_, we can maximize the net return by not treating any fields (*θ* = 0). If, on the contrary, *α < ϵ*_*w*_, then we can maximize the net return by treating a fraction of fields that corresponds to the critical value (*θ* = *θ*_*cw*_, which constitutes a high fraction of fields; Fig. 3B). We investigate *g*(*θ*) in more detail in Subsec. B.2.1 in S1 Appendix, where we show that the net return can either increase/decrease monotonically or exhibit a minimum as a function of *θ*.

Fig. 3C shows that the optimal coverage, *θ*^∗^, depends strongly on both the relative fungicide price, *f*, and the pathogen’s basic reproduction number, *R*_0_. This dependency exhibits three regimes for low, intermediate and high values of *R*_0_. For low *R*_0_ *<* 1, the optimal coverage does not depend on fungicide price because epidemics will not sustain even without treatment, and hence, not treating any field is optimal for any *f*. When *R*_0_ *>* 1, the optimal coverage does depend on *f*, showing a threshold pattern. For low *f* -values below the threshold, it is optimal to treat at least some of the fields (*θ*^∗^ *>* 0). For high *f* -values above the threshold, not treating any fields yields a maximum net return (*θ*^∗^ = 0). For intermediate *R*_0_-values, the threshold in *f* does not depend on *R*_0_ (hence, the boundary between white/gray and black areas is a vertical line in Fig. 3C). For high *R*_0_ values, the threshold in *f* becomes lower when *R*_0_ is increased. This threshold pattern, rather than a gradual increase in the optimal coverage with increasing *f*, occurs because, as we have shown above, the law of diminishing returns no longer holds when we increase the treatment coverage in multiple fields across a landscape. The identification of the threshold pattern can inform economic policy for achieving more sustainable fungicide use, as we elaborate in the Discussion.

Thus, the net return is maximized at extreme *θ*-values, and which of the extrema are optimal is determined by how the cost-ratio *α* relates to a combination of epidemiological parameters of the system.

#### Optimal fungicide coverage with resistance

Now, we focus on the fraction of fields to be treated with fungicide to maximize the net return in a landscape model when resistance exists. We consider a high-efficacy fungicide (*ϵ*_*w*_ = 1), no fitness cost of resistance (*β*_*r*_ = *β*_*w*_), and focus on partial resistance (0 *< ϵ*_*r*_ *<* 1) (fitness cost of resistance and full resistance are considered in Subsec. B.2.2 in S1 Appendix). The fungicide can still suppress the partially resistant strain with efficacy *ϵ*_*r*_, smaller than the efficacy against the wildtype strain, *ϵ*_*w*_. As before, the optimal fungicide coverage depends on the cost-ratio *α* (in Eq. (15)).

When *θ*_*cr*_ *>* 1 (*ϵ*_*r*_ *< ϵ*_*rc*_, where *ϵ*_*cr*_ is given by Eq. (11)), fungicide treatment is ineffectual in clearing the resistance. In this case, the optimal coverage, *θ*^∗^, depends on how the cost-ratio *α* relates to the combination of epidemiological parameters,

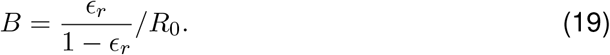

The expressions for *B* is similar to *A* in Eq. (18), but with *ϵ*_*r*_ instead of *ϵ*_*w*_. When a fungicide is sufficiently cheap, *α < B*, treating all fields maximizes the net return (*θ*^∗^ = 1; Fig. 4A). When a fungicide is too expensive, *α > B*, then not applying it is optimal (*θ*^∗^ = 0; Fig. 4A).

**Figure 4:**
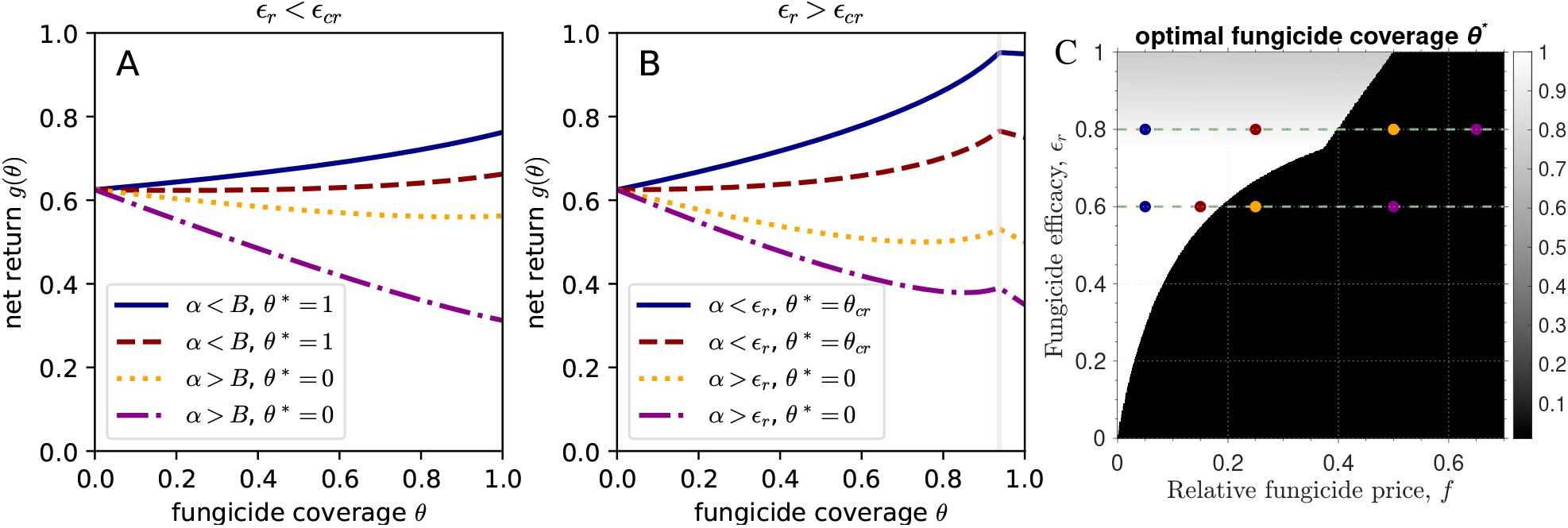
What is the optimal fungicide coverage in the landscape model with fungicide resistance? (A) The net return *g*(*θ*) is plotted against the fungicide coverage (fraction of treated fields *θ*) for a high degree of partial resistance (*ϵ*_*r*_ = 0.6 *< ϵ*_*cr*_ = 0.75). The four curves represent different relative fungicide prices (*f*): 0.05 (solid blue), 0.15 (dashed red), 0.25 (dotted orange), 0.5 (dashdotted magenta). (B) The net return *g*(*θ*) is plotted against *θ* for a low degree of partial resistance (*ϵ*_*r*_ = 0.8 *> ϵ*_*cr*_) which means that it is possible to drive the resistant strain to extinction by treating a high enough fraction of fields (*θ* = *θ*_*cr*_, vertical grey line). The four curves represent different values of *f* : 0.05 (solid blue), 0.25 (dashed red), 0.5 (dotted orange), 0.65 (dash-dotted magenta). (C) Heatmap of the optimal fungicide coverage *θ*^∗^ for varying fungicide price *f* and fungicide efficacy against resistant strain *ϵ*_*r*_. In the white region, it is optimal to treat all fields with fungicide; in the black region, it is optimal to not treat any fields; in the grey region, it is optimal to treat only a fraction of the fields. Each curve in (A) and (B) corresponds to a point on the green dashed line in (C). These points are highlighted by circles of the same color as lines in (A) and (B). Specifically, points on green dashed lines for *ϵ*_*r*_ = 0.6 corresponds to (A) and for *ϵ*_*r*_ = 0.8 corresponds to (B). Other parameter values are constant for all panels: *β* = 0.02, *R*_0_ = *βN/µ* = 4; *N* = 1000, *µ* = 5, *ϵ*_*w*_ = 1, *y* = 0.5.

When *θ*_*cr*_ *<* 1 (*ϵ*_*r*_ *> ϵ*_*cr*_), treating a sufficiently high fraction of fields (*θ > θ*_*cr*_) will drive the dominant resistant strain to extinction. Here, the optimal *θ* depends on how the cost-ratio *α* relates to the efficacy of the fungicide against the resistant strain *ϵ*_*r*_. If *α < ϵ*_*r*_, then treating a significant fraction of fields maximizes the net return (*θ*^∗^ = *θ*_*cr*_; Fig. 4B). If *α > ϵ*_*r*_, then not treating any field maximizes the net return (*θ* = 0; Fig. 4B). Fig. 4C provides a broader overview of the parameter space, considering how the optimal fungicide coverage *θ*^∗^ depends on fungicide price, *f*, and fungicide efficacy against the resistant strain, *ϵ*_*r*_.

Thus, when resistance is absolute, it is impossible to manage it apart from stopping or drastically reducing fungicide treatment (see Subsec. B.2.2 in S1 Appendix). Nevertheless, when resistance is partial, the net return can still be maximized by treating all fields or a significant fraction of fields (if the fungicide is sufficiently cheap) or by not treating any fields (if the fungicide is sufficiently expensive).

### 3.2 The economic cost of fungicide resistance

The maximum (or optimal) net return of a landscape becomes lower in the presence of resistance (compare Fig. 3A and Fig. 4A). Thus, the evolution of fungicide resistance can incur a substantial economic cost. Here, we propose a novel way to calculate the economic cost of fungicide resistance across a cultivated landscape using the expression for the net return in Eq. (14). Firstly, we compute the net return in the absence of resistance and in the presence of resistance according to Eq. (14). Secondly, we determine the optimal fraction of treated fields to maximize the net return in each of the cases. Finally, we calculate the economic cost of resistance, *C*_*R*_, as the difference between the optimum net return in the absence and in the presence of resistance:

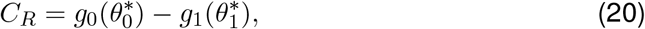

where 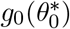 is the optimum net return and 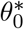 is the optimum coverage in the absence of resistance; 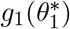 is the optimum net return and 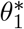 is the optimum coverage in the presence of resistance. In this calculation, we assume that, both with and without resistance, the optimum fraction of treated fields is known to farmers across the landscape and that they use these optimum values.

To illustrate this calculation, we plot the maximal net returns, 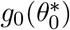 and 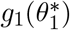, the optimal fractions of treated fields, 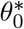 and 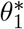, and the economic cost of resistance, *C*_*R*_, versus the relative fungicide price, *f* (Fig. 5). Fig. 5A compares the optimal net return in the absence and presence of resistance. The difference between these curves constitutes the economic cost of resistance evolution plotted in Fig. 5C. Next, we elucidate the dependence of the cost of resistance on the fungicide price *f*, the degree of resistance 1 − *ϵ*_*r*_, the basic reproduction number *R*_0_ and the yield loss 1 − *y*.

**Figure 5:**
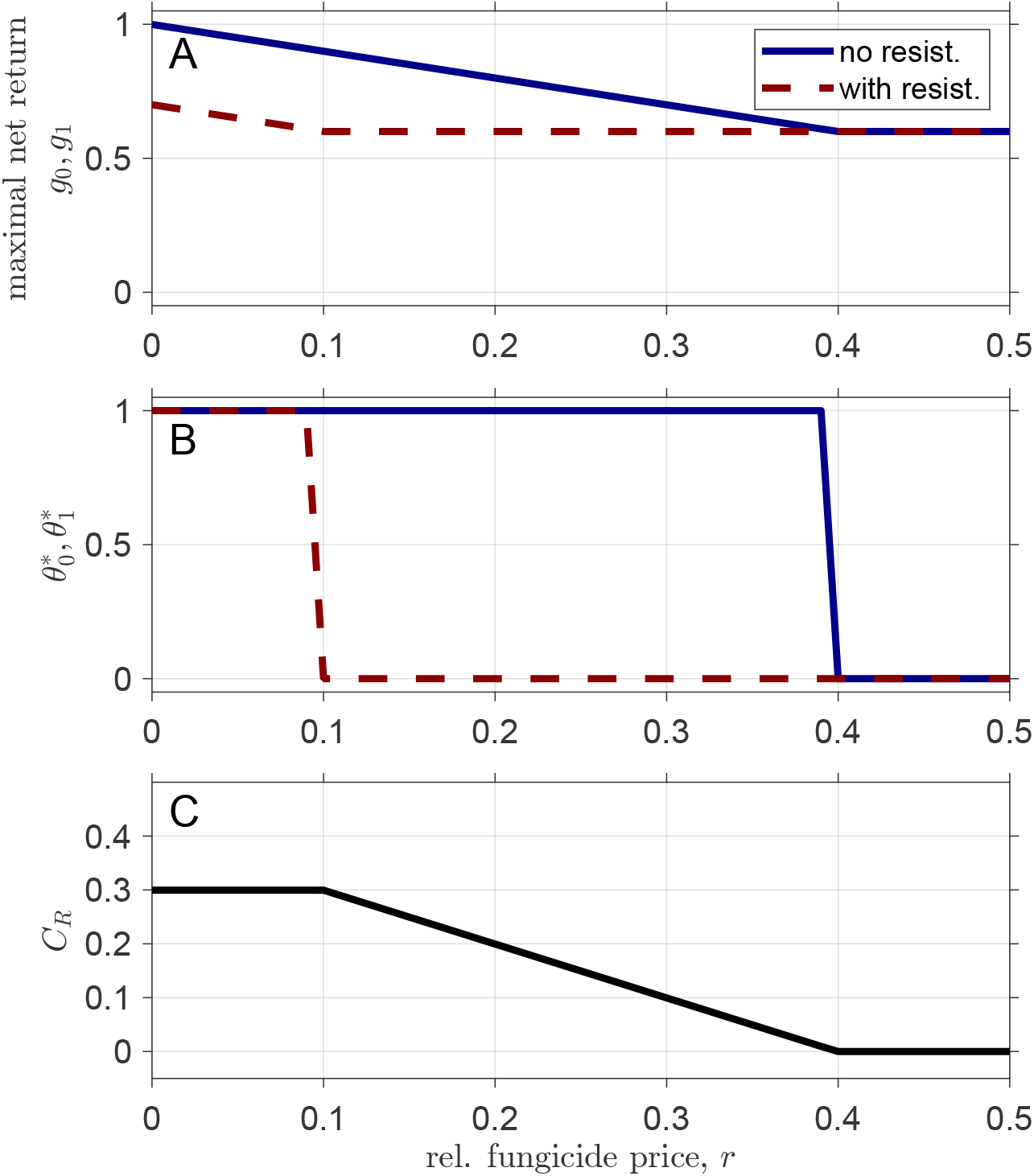
Calculation of the economic cost of fungicide resistance, *C*_*R*_, versus relative fungicide price, *f*, according to Eq. (20). (A) Optimal net return without resistance (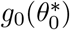, solid, blue curve) and with resistance 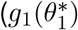, dashed, red curve). (B) Optimal fraction of treated fields without resistance (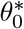, solid, blue curve) and with resistance (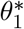, dashed, red curve). (C) Economic cost of resistance, *C*_*R*_, computed as the optimal net return without resistance minus the optimal net return with resistance. Parameter values: *β*_*r*_ = *β*_*w*_ = 0.025, *N* = 1000, *µ* = 5 (hence, *R*_0_ = *βN/µ* = 5); *ϵ*_*w*_ = 0.8, *ϵ*_*r*_ = 0.5.

#### Dependence on the fungicide price

Fig. 5C shows a typical dependence of *C*_*R*_ on *f* when the basic reproduction number of the pathogen exceeds its critical values (*R*_0_ *>* 1*/*(1 − *ϵ*_*w*_) in Eq. (6) and *R*_0_ *>* 1*/*(1 − *ϵ*_*r*_) in Eq. (12)). We explore three ranges of fungicide prices (*f*) – cheap, intermediate and expensive – corresponding to qualitatively different patterns in the *C*_*R*_ dependence. For cheap fungicides (low *f* -values), *C*_*R*_ remains constant and treating all fields maximizes the net return (Fig. 5B). In this range, the optimal net return drops linearly with increasing *f* at the same rate with and without resistance. Hence, the difference between the optimal net return with and without resistance remains constant (the economic cost of resistance in this parameter range is given by Eq. (C1) in S1 Appendix). For intermediate *f* -values, treating all fields remains optimal in the absence of resistance, but in the presence of resistance, the optimum switches to not treating any fields. The difference between the optimal net return without resistance and with resistance (that constitutes *C*_*R*_) declines linearly with *f* (the economic cost of resistance in this range is given Eq. (C2) in S1 Appendix). For expensive fungicides (high *f* -values), not treating any fields becomes optimal in the absence and in the presence of resistance. In this range, the cost of fungicide resistance remains at zero (Fig. 5C). Here, the optimal net return is determined by the yield of untreated fields, independent of the fungicide price *f* and irrespective of resistance.

In a different scenario, when *R*_0_ does not exceed the critical value for the wildtype strain [i.e., *R*_0_ *< R*_0*cw*_, Eq.(6)], *C*_*R*_ increases, reaches a maximum and then decreases as a function of *f*, reaching zero (region of the plot corresponding to 1 *< R*_0_ *<* 5 in Fig. 6B). This pattern arises due to a complex interplay between the optimal net return with and without resistance and the associated optimal fungicide coverage (we describe this in detail in Sec. C in S1 Appendix).

**Figure 6:**
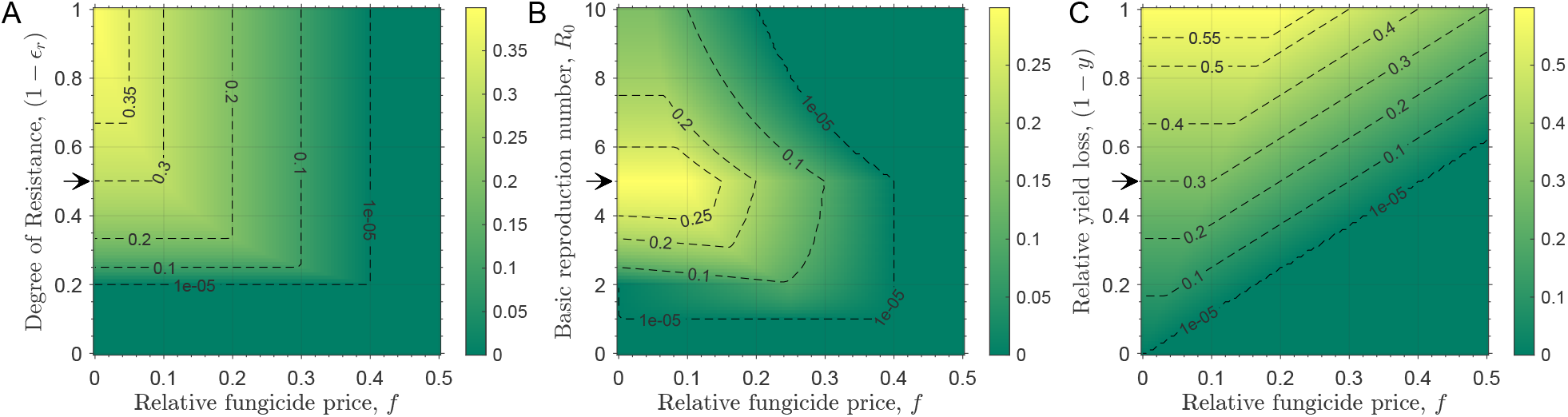
How does the economic cost of fungicide resistance, *C*_*R*_, depend on fungicide price (*f*), degree of resistance (1 − *ϵ*_*r*_), basic reproduction number (*R*_0_), and yield loss due to disease (1 − *y*)? Each panel illustrates how *C*_*R*_ depends on *f* along the *x*-axis and one of the other variables along the *y*-axis: (A) the effect of the degree of resistance, (1 − *ϵ*_*r*_), with *R*_0_ = 5 and *y* = 0.5; (B) the effect of the basic reproduction number, *R*_0_, with *ϵ*_*r*_ = 0.5 and *y* = 0.5 (*R*_0*cw*_ = 5, *R*_0*cr*_ = 2); (C) the effect of the yield loss due to disease, 1 − *y*, with *ϵ*_*r*_ = 0.5 and *R*_0_ = 5. Values of *ϵ*_*r*_, *R*_0_ and *y* used to plot Fig. 5 are highlighted with an arrow on the *y*-axis of panels (A), (B) and (C), respectively. Note that each panel uses a different grayscale mapping for *C*_*R*_ as shown in the grayscale bars. Parameter values that remain the same across all panels are: *N* = 1000, *µ* = 5, and *ϵ*_*w*_ = 0.8.

#### Dependence on the degree of resistance

The cost of resistance varies with the degree of resistance only for cheap fungicides (low *f* -values, Fig. 6A). Herein, *C*_*R*_ remains zero at low degrees of resistance, 1 − *ϵ*_*r*_, and then increases linearly with increasing 1 − *ϵ*_*r*_. Thus, as expected, higher degrees of resistance correspond to higher economic costs of resistance (compare horizontal parts of contour lines in Fig. 6A). This is because stronger resistance leads to higher numbers of fields becoming diseased despite treatment, which in turn results in lower overall yields of the landscape in the presence of resistance compared to the landscape in the absence of resistance.

For expensive fungicides (high *f* -values), the degree of resistance no longer affects the economic cost of resistance, which remains at zero in this range. In this scenario, the fungicide is so expensive that treatment is not economically justified, irrespective of resistance.

#### Dependence on the basic reproduction number

The dependence of the economic cost of resistance, *C*_*R*_, on the basic reproduction number, *R*_0_, is counter-intuitive (Fig. 6B). When the fungicide is expensive (the rightmost region in Fig. 6B), optimal fungicide coverage goes to zero both with and without resistance, and resistance incurs no economic cost. However, in a more interesting regime of low-to intermediately-priced fungicide, *C*_*R*_ exhibits complex, non-monotonic patterns versus *R*_0_ and *f* (Fig. 6B).

For cheap fungicides (i.e., small *f*), as *R*_0_ increases, *C*_*R*_ first remains at zero, then increases with *R*_0_, reaches a maximum and then decreases (compare different contour lines in Fig. 6B for small *f* -values). When *R*_0_ *<* 1, no epidemic occurs, and hence, the relative net return is maximal *g*_max_ = 1 both with and without resistance, resulting in *C*_*R*_ = 0. When 1 *< R*_0_ *< R*_0*cr*_ (see Eq. (12)), an epidemic can occur in both cases, but treating an optimal intermediate fraction of fields with a fungicide drives the pathogen to extinction: *θ*^∗^ = *θ*_*cw*_ without resistance and *θ*^∗^ = *θ*_*cr*_ with resistance. Hence, the optimal net return is given by *g*_max_ = 1 − *fθ*_*cr*_ with resistance and *g*_max_ = 1 − *fθ*_*cw*_ without resistance. Their difference *C*_*R*_ = *f* (*θ*_*cr*_ − *θ*_*cw*_), increases with *R*_0_ since *θ*_*cr*_ − *θ*_*cw*_ ∝ 1 − 1*/R*_0_ (see Eq. (5) and Eq. (10)). In other words, higher *R*_0_ values require increased fungicide coverage to eliminate the pathogen in both scenarios. However, due to lowered efficacy against the resistant strain, this coverage, and thus *C*_*R*_, increases faster in the presence of resistance.

For *R*_0*cr*_ *< R*_0_ *< R*_0*cw*_, applying fungicide to an optimal intermediate proportion of fields still drives the pathogen to extinction without resistance. While in the presence of resistance, treating all fields is optimal (*θ*^∗^ = 1), yet insufficient for elimination. In this regime, *g*_max_ without resistance continues to decline with *R*_0_ due to increasing required coverage. With resistance, *g*_max_ declines more rapidly due to both lower fungicide efficacy and higher prevalence of disease. As a result, *C*_*R*_ continues to increase with *R*_0_, reaching a maximum at *R*_0_ = *R*_0*cw*_.

Finally, and unexpectedly, when *R*_0_ *> R*_0*cw*_, *C*_*R*_ drops with increasing *R*_0_. In this case, fungicide treatment cannot extinguish the pathogen population, irrespective of the presence of resistance. To clarify, consider a simplified scenario of a cheap fungicide (*f* → 0). In this case, treating all fields (*θ*^∗^ = 1) becomes optimal both with and without resistance (Fig. 5B). Both with and without resistance, the optimal net return declines with *R*_0_ as the fraction of diseased fields increases with *R*_0_. Without resistance, this fraction (at endemic equilibrium) is given by *i*_*w*_ = 1 − 1*/* [*R*_0_(1 − *ϵ*_*w*_)]. With resistance, this fraction (also at endemic equilibrium, but now with the resistant strain) is given by *i*_*r*_ = 1 − 1*/* [*R*_0_(1 − *ϵ*_*r*_)]. *C*_*R*_ is proportional to the difference *i*_*r*_ − *i*_*w*_. Both *i*_*r*_ and *i*_*w*_ increase in a saturating manner with *R*_0_ (eventually tending to one at high *R*_0_-values) but do so at different, nonlinear rates in such a way that the difference between the two diminishes at higher *R*_0_-values, and so does the economic cost of resistance, *C*_*R*_. In other words, resistance incurs a lower economic cost at higher *R*_0_ values because even without resistance, it becomes challenging to suppress the disease due to the high invasiveness of the pathogen (as quantified by the basic reproduction number).

#### Dependence on yield loss

The economic cost of fungicide resistance *C*_*R*_ is influenced by the yield loss in diseased fields, 1 − *y* (relative to the yield of healthy fields; Fig. 6C). As intuitively expected, *C*_*R*_ increases with higher yield losses. But, surprisingly, for higher fungicide prices, this increase starts at higher values of yield loss. Thus, an estimation of the economic cost of fungicide resistance requires detailed knowledge of key epidemiological and economic parameters.

## 4 Discussion

Farmers can maximize the net economic return of fungicide application by optimizing the amount of fungicide applied based on the balance between the yield benefit and the economic costs of fungicide application (Jørgensen et al., 2017; Hillebrandt, 1960). However, this analysis neglects the indirect costs of fungicide application incurred by the evolution of fungicide resistance, rendering decisions based solely on this analysis unsustainable. One way to address this issue is to design economic policy instruments that incorporate indirect costs into fungicide prices, such as pesticide taxes or subsidies encouraging a reduction in pesticide application (Finger et al., 2017; Lee et al., 2019). However, this approach requires a robust estimation of the indirect costs, and a conceptual framework for such an estimation is absent from the existing literature. We build such a framework based on a simple but generic epidemiological model of crop epidemics across multiple fields combined with an economic analysis (Fig. 1). Using this framework, we demonstrated how to estimate the economic cost of the evolution of fungicide resistance in crop pathogens with long-distance dispersal.

Derivation of the economic cost of fungicide resistance evolution (*C*_*R*_) is a complex, multifaceted problem. However, with well-motivated assumptions, we derived analytical expressions for *C*_*R*_. This derivation allows us to investigate how *C*_*R*_ depends on the system’s critical epidemiological and economic parameters across entire ranges of plausible values. We found that *C*_*R*_ depends strongly on fungicide price, degree of resistance, pathogen’s basic reproduction number, and yield loss due to disease. As intuitively expected, *C*_*R*_ increases with the degree of resistance and yield loss due to disease. Unexpectedly, it remains constant or decreases with the fungicide price. Also unexpectedly, *C*_*R*_ shows a non-monotonic pattern as a function of the basic reproduction number, *R*_0_ (exhibiting a maximum at a critical value *R*_0_ = *R*_0*cw*_).

The three epidemiological parameters (degree of resistance, *R*_0_, and yield loss due to disease) can vary significantly across different crop diseases (Lucas et al., 2015; Mikaberidze et al., 2016; de Vallavieille-Pope et al., 2000; Savary et al., 2012). Even for a single disease, the three epidemiological parameters can vary substantially depending on environmental conditions, crop varieties and genetic composition of pathogen populations. A strong dependence of *C*_*R*_ on these parameters highlights the need for a robust empirical estimation of these parameters. Even after the estimation of the three parameters, designing economic policy instruments that work across different regions and crops would be demanding. These instruments are more likely to succeed if they are adjusted for specific regions and crops. These instruments also need to be adapted over time according to changes in cropping systems and pathogen populations. Implementing economic and ecological interventions in a realistic system should feed back into the parameter values. Thus, like adaptive therapy in medicine, the interventions are more likely to succeed if they are adapted to specific conditions and responsive in real-time.

One may consider the range of low *R*_0_-values (0 *< R*_0_ *< R*_0*cw*_) to be unrealistic for most fungal pathogens of crop plants. Firstly, according to empirical knowledge accumulated in plant pathology, fungal crop pathogens are challenging to eradicate (Lucas, 2020), suggesting that their *R*_0_ values should be much higher than one. Secondly, existing estimates of the basic reproduction number, e.g., for the major fungal pathogens of wheat *Zymoseptoria tritici* and *Puccinia striiformis* tend to be relatively high: in the range 4 − 10 for *Z. tritici* (Mikaberidze et al., 2017) and 20 − 30 for *P. striiformis* (Mikaberidze et al., 2016). The two arguments indicate that *R*_0_-values for major fungal crop pathogens will likely exceed the eradication thresholds. Accordingly, it would make sense to focus more on the range of higher *R*_0_-values, where it exceeds the eradication thresholds (*R*_0_ *> R*_0*cw*_, *R*_0_ *> R*_0*cr*_). At the same time, here we used *R*_0_ as determined at the landscape scale, while most experimental estimates for fungal pathogens have been evaluated at the individual field scale. Due to lower transmission rates and greater spatial regularity in individual susceptible units, we expect lower *R*_0_ estimates at the landscape scale than at the field scale, at least for certain landscape configurations (Mikaberidze et al., 2016; van den Bosch et al., 2024). In this case, the estimation of *C*_*R*_ across the entire range of *R*_0_ values, as we did here, becomes of interest.

Along the way to calculating the economic cost of fungicide resistance, we also found that the law of diminishing returns does not hold for the economic return from fungicide applications over a cultivated landscape (Fig. 2D). Optimal fungicide dosing for individual fields still exhibits the law of diminishing returns for the net economic return (Fig. 2B). Hence, the economically optimum dose decreases gradually with increasing fungicide price. However, for multiple fields across the landscape, the optimal proportion of treated fields also decreases with the ratio of fungicide price and yield loss (*α*) but does so via a sudden jump (see Fig. 5B). Thus, adjusting the magnitude of fungicide treatment to maximize the net return is more difficult at the landscape scale owing to coordinating fungicide applications across multiple farms and handling the discontinuous nature of how *θ*^∗^ depends on *α*. At the same time, this presents an opportunity for economic policy to minimize pesticide use because a limited increase in *α* induced by a policy instrument may lead to a disproportionately high, sudden drop in the optimal fungicide coverage, *θ*^∗^. However, achieving this optimum requires cooperation between farmers across a regional landscape.

Our study considers a binary choice in fungicide application. In reality, farmers can adjust the fungicide dose (Hobbelen et al., 2014; Mikaberidze et al., 2017), use mixtures or alternations of different fungicides (van den Bosch et al., 2014; Mikaberidze et al., 2014). Furthermore, the farmers can adopt disease-resistant crop varieties (Carolan et al., 2017; Taylor and Cunniffe, 2023) and cultural control measures (such as crop rotations (Shah et al., 2021; Bargués-Ribera and Gokhale, 2020), cultivar mixtures (Finckh et al., 2000; Mikaberidze et al., 2015; M’Barek et al., 2020) and restrictions on growing certain crops, e.g., soybean-free periods in Brazil (Godoy et al., 2016)). These scenarios can be incorporated into our modelling framework to explore how different control measures can protect each other from pathogen adaptation (Carolan et al., 2017; Kristoffersen et al., 2020; Corkley et al., 2025) and mitigate the economic cost of pathogen adaptation.

We calculated the economic cost of fungicide resistance as a benchmark corresponding to an optimistic scenario where all farmers across the landscape coordinate their fungicide applications to reach the optimum proportion of treated fields that gives the maximum net return for the entire landscape. However, the choices made at individual farms do not always align with the landscape-wide optimum. Instead, they are influenced by access to information, risk perceptions, socio-demographics, structure of farm business, and other factors (Edwards-Jones, 2006; Murray-Watson et al., 2022; Olita et al., 2024; Otoo et al., 2024). To address this, models should incorporate farmers’ decision-making heuristics and dynamic choices (Gokhale and Sharma, 2023). This would allow us to adjust the fungicide coverage and explore how close to the landscape-scale optimum one can get using different decision-making approaches. Including a socioeconomic point of view brings us closer to identifying policy interventions most likely to maximize crop protection’s economic return at a regional scale.

The methodological novelty of our study lies in extending a spatially implicit metapopulation model of plant disease epidemics (Parnell et al., 2006) to incorporate an economic analysis of fungicide use. Landscape perspective is recognized as crucial in plant disease epidemiology (Plantegenest et al., 2007), and more recently, spatially explicit landscape-scale models addressed a range of epidemiological/evolutionary questions focusing mainly on pathogen adaptation to disease resistance in plants (Fabre et al., 2012; Papaïx et al., 2014; Papaïx et al., 2015; Djidjou-Demasse et al., 2017; Rimbaud et al., 2021). While some studies recognized the importance of the economic aspects of plant disease epidemics, they did not incorporate an explicit economic analysis. Qualitative conceptual frameworks integrating plant pathogen adaptation to control measures with socioeconomic aspects are only starting to appear (Geffersa et al., 2023) and remain a mathematical challenge (Cunniffe et al., 2014).

Extending the marginal theory of pesticide use (Hillebrandt, 1960; Jørgensen et al., 2017) from the scale of individual farms to a broader scale of cultivated landscapes, we incorporated economic analysis in our epidemiological study. More specifically, we extended the concept of economic yield (Zadoks and Schein, 1979; Nutter et al., 1993) to the landscape scale. Earlier, Te Beest et al. (2013) extended the marginal theory of pesticide use by incorporating the stochasticity of plant disease epidemics in calculating the optimal fungicide dose, maximizing the net return from a fungicide application programme at the scale of an individual farm. This approach has been subsequently developed to incorporate the evolution of fungicide resistance (van den Bosch et al., 2018, 2020). Nevertheless, focusing on individual farm management, they neglected the externalities of the evolution of resistance. Here, we addressed this issue by extending the marginal theory of pesticide use (Hillebrandt, 1960; Jørgensen et al., 2017) from individual farms to cultivated landscapes in a mathematical approach that incorporates the epidemic development and the evolution of fungicide resistance and links these processes with an economic analysis. Given that our framework builds on fundamental equations of epidemiology and economic cost-benefit analysis, we envision applying our framework to modelling other biotic stresses of crops, such as insect pests and weeds (Lauenroth and Gokhale, 2023).

## 5 Conclusion

We have developed a bioeconomic framework combining a plant disease epidemiological model with an economic analysis at the landscape scale. Using this approach, we found that the law of diminishing returns does not hold for economic returns from fungicide applications across cultivated landscapes. This is surprising given the almost universal status of this law. The breakdown of this law suggests that adjusting the amount of fungicide applied to maximize the net return is challenging at the landscape scale. The intervention needs to be coordinated across multiple farms, and the discontinuous nature of how the optimal coverage depends on fungicide price requires a drastic change in fungicide coverage.

Using the bioeconomic modeling framework, we calculated the economic cost of the evolution of fungicide resistance, *C*_*R*_, which constitutes our study’s primary novel outcome. We found that *C*_*R*_ depends strongly on the fungicide price, the basic reproduction number of the pathogen, its degree of fungicide resistance and the consequential yield loss. While intuitively, *C*_*R*_ increases with the degree of resistance and yield loss due to disease, surprisingly, it declines with the fungicide price and exhibits a complex non-monotonic pattern as a function of the basic reproduction number. Hence, to estimate *C*_*R*_, robust estimations of these parameters are necessary, incorporating environmental variation, crop varieties and genetic composition of the pathogen population. Estimating the economic cost of resistance would then inform economic policy instruments to encourage more sustainable fungicide use.

Moving forward, our framework presents an opportunity for future empirical studies to validate the findings presented here across various agricultural landscapes and pathogen-crop systems. Future research focusing on extending our modeling framework to incorporate spatial heterogeneity, stochasticity, and multiple pathogen interactions will improve its applicability and help refine the estimation of the economic cost of fungicide resistance. Furthermore, incorporating farmers’ decision-making heuristics and economic policy tools, such as taxes and/or subsidies, into our bioeconomic modelling framework may help in designing more effective policy interventions for sustainable disease management. Integrating our framework with real-time monitoring and predictive modeling of crop epidemics will have the potential to improve disease management. Ultimately, we envision this study to provide a conceptual and methodological basis for a fruitful integration of economics and eco-evolutionary dynamics necessary for achieving more sustainable agriculture.

## Supporting information

**S1 Appendix. Supplemental methods and analysis: Stability, Optimization and Economic Cost of Resistance**. (A) Linear stability analysis and computation of fixed points and basic reproduction number. (B) Maximizing economic return of fungicide treatments. (C) The economic cost of fungicide resistance.

## Acknowledgments

C.S.G. acknowledges funding from the Max Planck Society. P.V. gratefully acknowledges support from the University of Oldenburg, University of Bielefeld, HIFMB and the University of California, Berkeley. A.M. gratefully acknowledges funding from the University of Reading (UK) and ETH Zurich (Switzerland).

## Data Availability

All data and codes used in this study are available on Zenodo (https://doi.org/10.5281/zenodo.15557902). Any necessary dependencies, including specific versions of software packages and libraries, instructions for using the codes, are also documented in the repository’s README file.

## Authors’ contributions

Conceptualization, A.M., C.G.S., M.B.R. and P.V.; Methodology, A.M., C.G.S., M.B.R. and P.V.; Investigation, A.M., C.G.S. and P.V.; Writing – Original Draft, A.M., C.G.S. and P.V.; Writing – Review & Editing, A.M., C.G.S., M.B.R. and P.V.; Visualization, A.M. and P.V.; Project administration, A.M., C.G.S.,and P.V.;

## Supporting information S1 Appendix

### A SI. Linear stability analysis and computation of fixed points and basic reproduction number

#### A.1 Model without fungicide treatment

We will first calculate the basic reproduction number for the case without fungicide treatment, i.e. *θ* = 0. In this case, we have only two state variables *H*_*u*_ and *I*_*uw*_, since *H*_*t*_ = 0 and *I*_*tw*_ = 0. The two fixed points for the set of ordinary differential equations (ODEs) Eq.(1) are 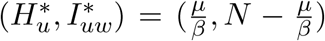 and (*N*, 0). Depending on the rate of transmission (*β*) and recovery (*µ*), the linear stability analysis of the system produces either of the two scenarios:

- (*N*, 0) is unstable and 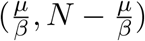 is stable when *βN > µ*;
- (*N*, 0) is stable and 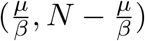 is unstable when *βN < µ*.

The Jacobian of the dynamical system is given by *J*(*I*_*uw*_) = *β*(*N* − 2*I*_*uw*_) − *µ*. Assuming, *H*_*u*_(0) ≈ *N* and *I*_*uw*_(0) ≈ 0, we obtain the following expression for the basic reproduction number using the Routh-Hurwitz criterion:

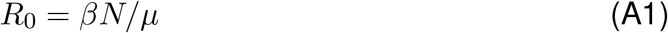

If *R*_0_ *>* 1, during the initial phase of the epidemic, when almost all the fields are healthy, there is an exponential growth in the number of untreated infected fields given by *I*_*uw*_(*t*) = *I*_*uw*_(0) exp[(*R*_0_ − 1)*µt*].

#### A.2 Model with fungicide treatment

##### A.2.1 Computation of fixed points and their stability for the treatment scenario

When we include fungicide treatment (*θ >* 0), the epidemiological dynamics of untreated and treated infected fields (*I*_*uw*_ and *I*_*tw*_) is given by:

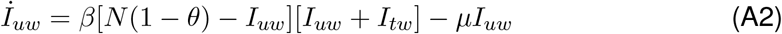

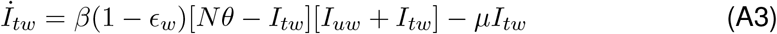

Here, we have used Eq. (2) to replace *H*_*u*_ and *H*_*t*_ from Eq. (A2)-(A3). In this case, there are three fixed points: FP1, FP2 and FP3. Fixed point 1 (FP1) corresponds to the disease-free equilibrium

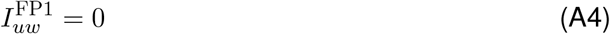

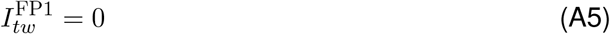

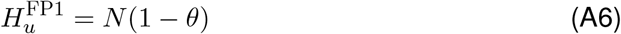

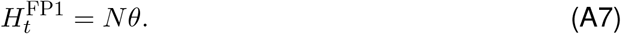

Fixed point 2 (FP2) corresponds to the endemic equilibrium

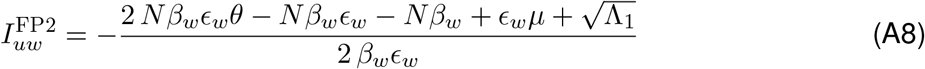

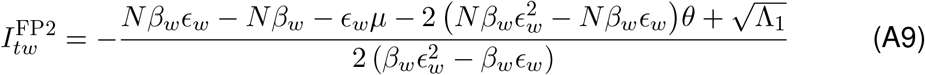

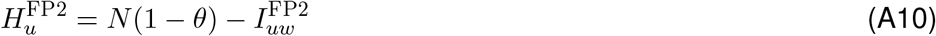

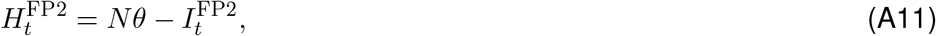

Where

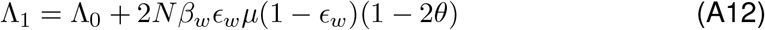

And

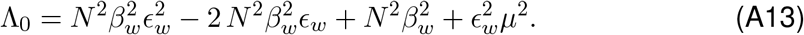

Fixed point 3 (FP3) also corresponds to non-trivial values of the four state variables

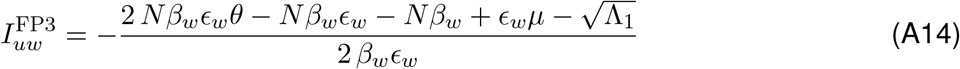

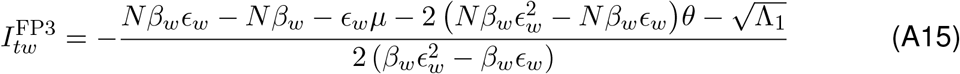

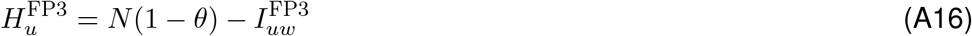

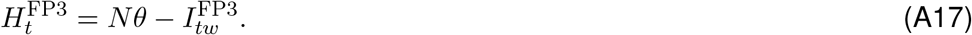

However, 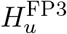and 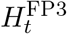 exhibit negative values across biologically plausible parameter regimes and therefore do not correspond to a biologically meaningful solution.

The three fixed points and their biological interpretations are summarized in Table S1. Note that the names of the fixed points in the Table S1 may not be strictly correct, as they depend on the stability conditions of the eigenvalues expressions derived using Matlab code in the following Github folder. The stability conditions and the expressions for the fixed points can also be found in Github folder.

**Table S1:**
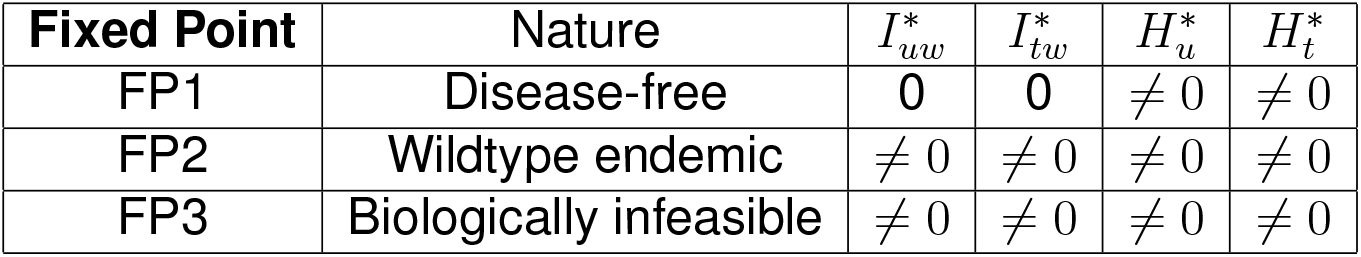
Fixed points for the treatment scenario.

| Fixed Point | Nature | $I_{uw}^*$ | $I_{tw}^*$ | $H_u^*$ | $H_t^*$ |
| --- | --- | --- | --- | --- | --- |
| FP1 | Disease-free | 0 | 0 | $\neq 0$ | $\neq 0$ |
| FP2 | Wildtype endemic | $\neq 0$ | $\neq 0$ | $\neq 0$ | $\neq 0$ |
| FP3 | Biologically infeasible | $\neq 0$ | $\neq 0$ | $\neq 0$ | $\neq 0$ |

##### A.2.2 Computation of basic reproduction number in the limiting case of a high-efficacy fungicide

We analyze the special case when the fungicide efficacy *ϵ*_*w*_ = 1, i.e. the treated fields are completely protected from infection. The above Eq. (A2)-(A3) then becomes,

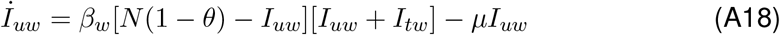

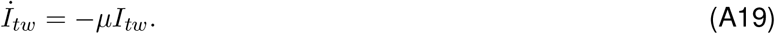

There are two fixed points of the system of (Eq. (A18)-(A19)) 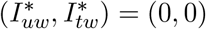 and 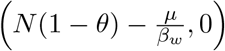. The Jacobian matrix of the above equation is 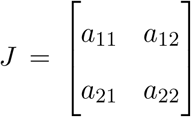 where the elements are given by:

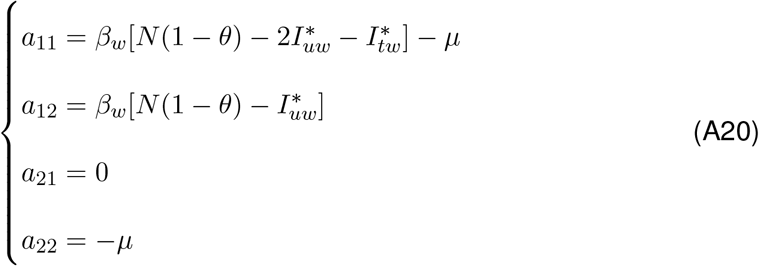

The stability of the fixed point can be evaluated from the trace tr(*J*) and the determinant det(*J*) of the Jacobian:

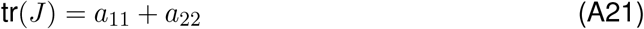

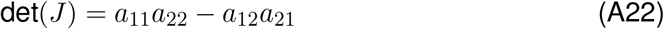

Let *J*_1_ and *J*_2_ be the Jacobian matrix of the of the two fixed points (0, 0) and 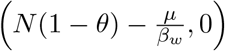 respectively. The fixed point 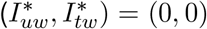 is stable if tr(*J*_1_) *<* 0 and det(*J*_1_) *>* 0, which happens for *Nβ*_*w*_(1 − *θ*) − *µ >* 0 or *R*_0_(1 − *θ*) *<* 1. Similarly, the fixed point 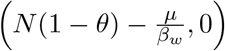 is stable when tr(*J*_2_) *<* 0 and det(*J*_2_) *>* 0 which happens for *Nβ*_*w*_(1 − *θ*) − *µ >* 0 or *R*_0_(1 − *θ*) *>* 1. Hence, the infected field can invade the population of healthy fields if *R*_0_(1 − *θ*) *>* 1 in the limiting case of a high efficacy fungicide *ϵ*_*w*_ = 1. Thus, ℛ_0_ = *R*_0_(1 − *θ*) is the effective reproduction number of the system in this case. Let us now derive the expression for ℛ_0_ for a general value of fungicide efficacy 0 ≤ *ϵ*_*w*_ ≤ 1 using the next-generation matrix method.

##### A.2.3 Computation of basic reproduction number for the general case of a partial-efficacy fungicide

Above, we derived the condition for the invasion of healthy fields by infected using stability criteria for the the Jacobian matrix. An alternative approach is to directly derive the reproductive number ℛ_0_ for our system by using the next generation matrix. The proof and detailed discussion of this method is given in Diekmann et al. (1990); van den Driessche and Watmough (2002). Here, we will follow the prescription outlined in van den Driessche (2017) to evaluate ℛ_0_.

Consider the equation, ^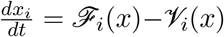^ for *i* = 1, 2, …,*m*where x = [*x1, x2*, …, *xn*] is the number of individuals in various compartments and *m < n* is the total number of infected compartments. Now if 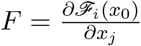 and 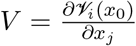 for 1 ≤ *i, j* ≤ *m* and *x*_0_ is the infection free state, then ℛ_0_ is the maximum real part of the eigenvalues of *FV* ^−1^.

When there are no infected fields, *x*_0_ = [*H*_*u*_, *H*_*t*_, *I*_*uw*_, *I*_*tw*_] = [*N* (1 − *θ*), *Nθ*, 0, 0], then

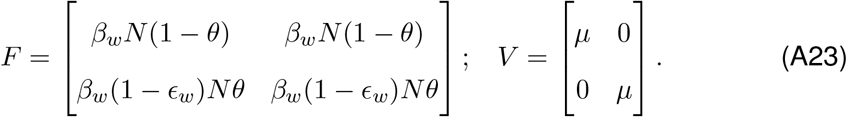

Hence *FV* ^−1^ is given by

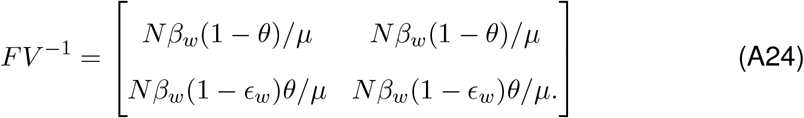

The ℛ_0_ is the root corresponding to the maximum real part of the characteristic equation,

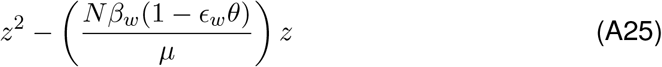

Hence, 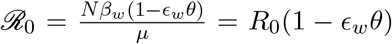 where *R*_0_ = *Nβ*_*w*_*/µ*. The expression for the basic reproduction number for the general case of a partial-efficacy fungicide has also been derived via Matlab code provided in the Github repository.

#### A.3 Model with fungicide treatment and resistance

##### A.3.1 Special case 1: Computation of fixed points and their stability for a high-efficacy fungicide and absolute resistance

The equations describing the epidemiological dynamics with fungicide resistance, using Eq. (7) and assuming *ϵ*_*w*_ = 1 and *ϵ*_*r*_ = 0, are given by:

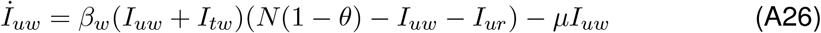

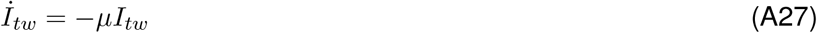

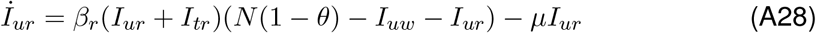

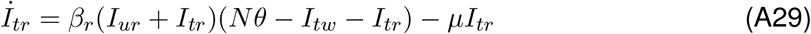

The fixed points can be categorized into four biologically significant solutions. The first solution corresponds to disease-free equilibrium, where no infected fields exist. The second solution corresponds to the wildtype endemic equilibrium, where healthy and wildtype infected fields co-exist. The third solution corresponds to the resistant endemic equilibrium, where healthy and resistant infected fields co-exist. Lastly, the fourth solution corresponds to coexistence between the healthy fields, wildtype infected and resistant infected fields. Fixed points and their biological interpretations are summarized in Table S2.

**Table S2:** Fixed points of the ODE system with fungicide resistance in the limiting case of *ϵ*_*w*_ = 1 and *ϵ*_*r*_ = 0 (Special Case 1).

| Fixed Point | Nature | $I_{uw}^*$ | $I_{tw}^*$ | $I_{ur}^*$ | $I_{tr}^*$ |
| --- | --- | --- | --- | --- | --- |
| FP1 | Disease-free | 0 | 0 | 0 | 0 |
| FP2 | Wildtype endemic | $\neq 0$ | 0 | 0 | 0 |
| FP3 | Resistant strain endemic | 0 | 0 | $\neq 0$ | $\neq 0$ |
| FP4 | Coexistence | $\neq 0$ | 0 | $\neq 0$ | $\neq 0$ |

Note that the names of the fixed points in Table S3 may not be strictly correct, as they depend on the stability conditions. Using Eq (A26)-(A29), we can compute four Jacobian matrices for the four fixed point conditions. We know that a fixed point is stable if all the real parts of eigenvalues of the corresponding Jacobian matrix are negative. Hence, we can derive the conditions on the parameters of the systems for all the fixed points to remain stable. The expressions for the stability conditions and the fixed points can be found in the following Github folder while the corresponding Matlab code can be found in the following Github folder.

##### A.3.2 Special case 2: Computation of fixed points and their stability for a high-efficacy fungicide and partial resistance

When the fungicide has the highest efficacy (*ϵ*_*w*_ = 1), but resistance is partial (0 *< ϵ*_*r*_ *<* 1), there are five fixed points summarized in Table S3. As before, the names of the fixed points provided in Table S3 may not be strictly correct, as they depend on the stability conditions. The non-zero expressions in Table S3 for 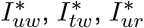 and 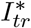, and the stability conditions of the fixed points are too long to be displayed in the text. Hence, we provide them in the following Github folder, while the corresponding Matlab code can be found in the this Github folder.

**Table S3:** Fixed points of the ODE system with partial fungicide resistance and *ϵ*_*w*_ = 1 (Special Case 2).

| Fixed Point | Nature | $I_{uw}^*$ | $I_{tw}^*$ | $I_{ur}^*$ | $I_{tr}^*$ |
| --- | --- | --- | --- | --- | --- |
| FP1 | Disease-free equilibrium | 0 | 0 | 0 | 0 |
| FP2 | Resistant strain endemic (unstable) | 0 | 0 | $\neq 0$ | $\neq 0$ |
| FP3 | Wild type strain endemic | $\neq 0$ | 0 | 0 | 0 |
| FP4 | Coexistence | $\neq 0$ | 0 | $\neq 0$ | $\neq 0$ |
| FP5 | Resistant strain endemic | 0 | 0 | $\neq 0$ | $\neq 0$ |

##### A.3.3 General Case: Computation of fixed points and their stability with fungicide resistance

When *ϵ*_*w*_ and *ϵ*_*r*_ have general values, there are six fixed points for the epidemiological dynamics described by Eq. (7). These fixed points are given in Table S4.

**Table S4:** Fixed Points of the ODE System with fungicide resistance (General Case)

| Fixed Point | Nature | $I_{uw}^*$ | $I_{tw}^*$ | $I_{ur}^*$ | $I_{tr}^*$ |
| --- | --- | --- | --- | --- | --- |
| FP1 | Disease-free equilibrium | 0 | 0 | 0 | 0 |
| FP2 | Coexistence | $\neq 0$ | $\neq 0$ | $\neq 0$ | $\neq 0$ |
| FP3 | Wildtype endemic | $\neq 0$ | $\neq 0$ | 0 | 0 |
| FP4 | Resistant endemic | 0 | 0 | $\neq 0$ | $\neq 0$ |
| FP5 | Resistant endemic | 0 | 0 | $\neq 0$ | $\neq 0$ |
| FP6 | Wildtype endemic | $\neq 0$ | $\neq 0$ | 0 | 0 |

Again, the names of the fixed points provided in Table S4 may not be strictly correct, as they depend on the stability conditions. The non-zero expressions in Table S4 for 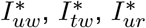 and 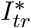 and the stability conditions of the fixeds point are too large to be displayed in the text, hence we provide them in the following Github folder, while the corresponding Matlab code can be found in this Github folder.

##### A.3.4 Computation of basic reproduction number in the general case with fungicide resistance

We follow the previous formulation of computing *R*_0_ via the next-generation matrix method for the case with fungicide resistance. The epidemiological dynamics are described by Eq. (7). Using Eq. (7), matrix *F* and *V* can be evaluated as:

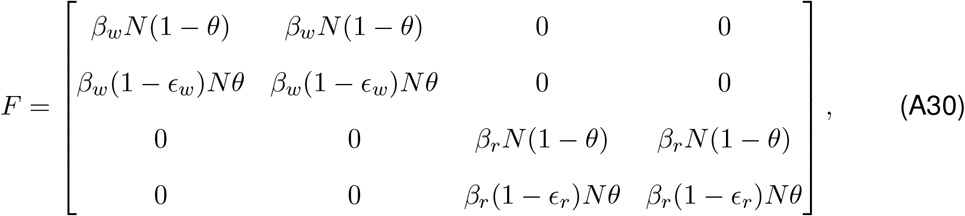

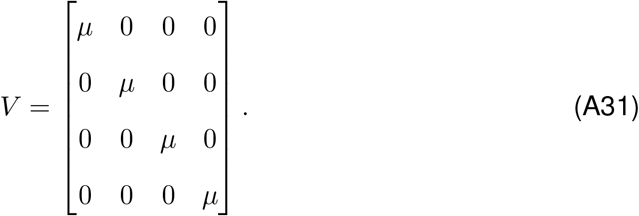

Here, we assume that initially there are no infected fields *x*_0_ = [*H*_*u*_, *H*_*t*_, *I*_*uw*_, *I*_*tw*_, *I*_*ur*_, *I*_*tr*_] = [*N* (1 − *θ*), *Nθ*, 0, 0, 0, 0]. *R*_0_ is the maximum real part of the eigenvalues of *FV* ^−1^. *FV* ^−1^ has two real valued eigenvalues *λ*_1_ = *Nβ*_*w*_(1−*ϵ*_*w*_*θ*)*/µ* and *λ*_2_ = *Nβ*_*r*_(1−*ϵ*_*r*_*θ*)*/µ*. Hence, *R*_0_ = max(*λ*_1_, *λ*_2_).

### B SI. Maximizing economic return of fungicide treatments

#### B.1 Optimal intensity of fungicide treatments in individual fields versus cultivated landscapes

How does the net economic return depend on the intensity of fungicide treatment in individual fields and in cultivated landscapes? To answer this question, we derive mathematical expressions for the net economic return as a function of the intensity of fungicide treatment in individual fields and in cultivated landscapes. We use these in Sec. Optimal fungicide coverage without resistance of the main text to contrast individual fields and cultivated landscapes.

##### B.1.1 OPTIMAL net return in an individual field

When managing disease in an individual field, growers can adjust the intensity of control by optimizing the fungicide dose. To determine the optimal dose, we consider the expression for the net return that is similar to Eq. (14) in the main text, but here we formulate it for the case of an individual field

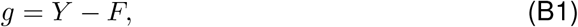

Where

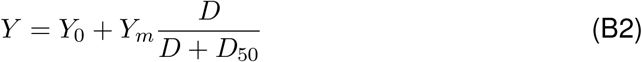

is the yield (expressed as the fraction of the maximum yield converted into its monetary equivalent) when a dose *D* of the fungicide has been applied. The first term in Eq. (B2) is the yield without fungicide application, *Y*_0_. The second term in Eq. (B2) is the yield benefit of the fungicide application, where the functional form used is a simplification of the Hill function, which describes well typical empirical dose-response relationships for fungal diseases in cereal crops (Mikaberidze et al., 2017). The parameter *Y*_*m*_ represents the maximum effect at large doses, and *D*_50_ corresponds to the dose for which half of the maximum effect is achieved. Hence, *Y*_0_ + *Y*_*m*_ corresponds to the attainable yield (Zadoks and Schein, 1979; Nutter et al., 1993).

The second term, *F*, in Eq. (B1) is the total fungicide price. We consider it to be proportional to the fungicide dose *D*

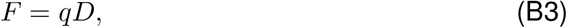

where *q* is the price per unit fungicide. To compute the net return in Eq. (B1), we measure *F* in the same units as *Y* : as the fraction of the maximum yield converted into its monetary equivalent. We substitute Eq. (B2) and Eq. (B3) into Eq. (B1) and obtain the following expression of the net return as a function of the fungicide dose (Eq. (16) in the main text)

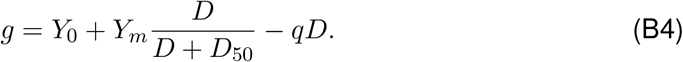

Net return, *g*, in Eq. (B4) exhibits a maximum at the fungicide dose

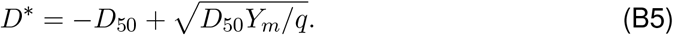

If the fungicide becomes sufficiently expensive (*q > Y*_*m*_*/D*_50_), the expression for the optimum dose in Eq. (B5) becomes negative, meaning that the net return is maximized when no fungicide is applied. For less expensive fungicides, the net return does exhibit a maximum at positive doses. The optimum dose is higher for cheaper fungicides; the optimum dose is also higher for fungicides with a higher maximum effect, *Y*_*m*_.

The yield that corresponds to the maximum net return is then given by

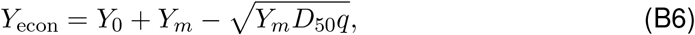

and is defined as the economic yield (Zadoks and Schein, 1979; Nutter et al., 1993). The economic yield decreases from the attainable yield, *Y*_0_ + *Y*_*m*_ (achieved if the fungicide is very cheap, i.e. *q* ≈ 0), by a value proportional to the square root of the fungicide price per unit *q*. We use the expression for the net return in Eq. (B4) above and its decomposition into *Y* and *F* to illustrate the law of diminishing returns in the context of an individual field in Fig. 2A,B of the main text.

##### B.1.2 Optimal net return in a cultivated landscape

As in the section above, we decompose the expression for the net return into the total yield, *Y* (*θ*), and the total fungicide price, *F* (*θ*), but now in the context of a cultivated landscape and hence both of these are functions of *θ*:

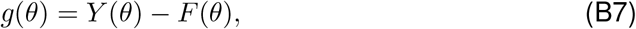

Where

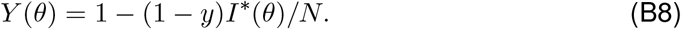

Here, *I*^∗^(*θ*) is number of diseased fields at a stable fixed point that can be computed from the model Eq. (7) in the main text.

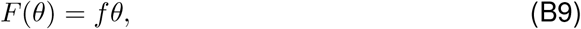

where *f* = *c/y*_*H*_ is the relative fungicide price. We use the expression for the net return in Eq. (B7) above and its decomposition into *Y* (*θ*) and *F* (*θ*) to illustrate the absence of diminishing returns when controlling disease across multiple fields in a landscape in Fig. 2C,D of the main text.

#### B.2 Analysis of the optimal fungicide coverage across landscapes

We investigate the scenario when the system has reached a stable fixed point and analyze the expression for the associated net return given by Eq. (14) in the main text

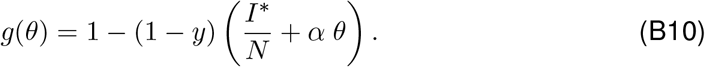

##### B.2.1 Optimal fungicide coverage without resistance

###### The case of high *R*_0_ when fungicide treatment cannot drive the pathogen population to extinction (*θ*_*cw*_ *>* 1)

In this regime, the fixed point that corresponds to the endemic equilibrium (FP2 in SI A.2.1) is stable and is therefore realized (over sufficiently large times) across the entire range of *θ*-values. The associated equilibrium values of the state variables are given by Eqs. (A8)-(A11) in SI A.2.1. We calculate the net return by substituting these values into Eq. (B10). The resulting dependence *g*(*θ*) is illustrated in Fig. 3A. The four curves in Fig. 3A differ only in the values of the cost-ratio *α* and represent four qualitatively different scenarios. The four curves start from the same value at *θ* = 0, because in the absence of fungicide application, the net return *g*(*θ*) in Eq. (B10) does not depend on the fungicide price *f*. Which of the four regimes is realized depends on how *α* relates to the three constants *A*_1_, *A*_2_ and *A*_3_ that are independent of *θ*, but do depend on the pathogen’s basic reproductive number in the absence of treatment *R*_0_ = *βN/µ* and on the fungicide efficacy *ϵ*_*w*_:

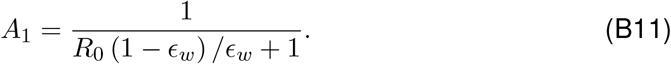

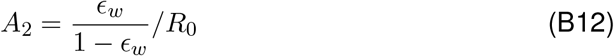

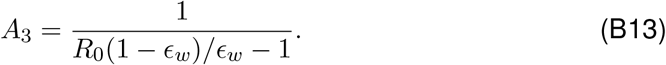

When *α < A*_1_, the net return grows monotonically as we treat more fields and thereby increase *θ* (see Fig. 3A). When *A*_2_ *< α < A*_1_, *g*(*θ*) exhibits a minimum at an intermediate value of *θ* (see Fig. 3A). If in addition *α < A*_3_ (i.e., *A*_2_ *< α < A*_3_), then *g*(*θ* = 1) *> g*(*θ* = 0), meaning that treating all fields gives a higher net return than no treating any fields at all; moreover *θ* = 1 gives the highest net return among all *θ*-values under this scenario (see Fig. 3A). Alternatively, if *A*_3_ *< α < A*_1_ the dependency *g*(*θ*) still exhibits a minimum at an intermediate value of *θ*, but now *g*(*θ* = 0) *> g*(*θ* = 1), meaning that not applying the fungicide in any of the fields gives a higher net return than applying the fungicides on all fields (see Fig. 3A). In this case, *θ* = 0 gives the highest net return among all *θ*-values. Finally, when *α > A*_1_, the net return *g*(*θ*) decreases monotonically with increasing *θ* and not treating any fields at all (i.e., *θ* = 0) maximizes the net return (see Fig. 3A). To summarize, when *θ*_*cw*_ *>* 1, to maximize the net return, one should either treat all fields (*θ*^∗^ = 1 if *α < A*_2_), or treat no fields (*θ*^∗^ = 0 if *α > A*_2_).

###### The case of low *R*_0_ when fungicide treatment can drive the pathogen population to extinction (*θ*_*cw*_ *<* 1)

Next, consider the regime when the critical fraction of treated fields in Eq. (5) is lower than one (*θ*_*cw*_ *<* 1). In this regime, two different fixed points can become stable and be realized (over sufficiently large times) depending on the values of *θ*. At *θ < θ*_*cw*_, the fixed point corresponding to the endemic equilibrium is stable (FP2), while at *θ > θ*_*cw*_, the fixed point corresponding to the disease-free equilibrium is stable. Also here, *g*(*θ*) exhibits four qualitatively different regimes depending on how the parameter *α* relates to the three constants 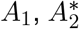 and 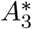. Here, *A*_1_ is given by Eq. (B11), 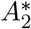 and 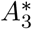 are given by

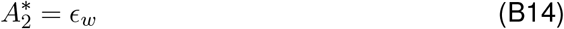

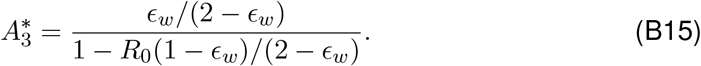

Here, 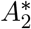 and 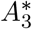 do not depend on *θ*, but do depend on *R*_0_ and *ϵ*_*w*_.

The net return versus the fungicide coverage *g*(*θ*) exhibits different patterns in the two different ranges of *θ*-values. When *θ < θ*_*cw*_, the behavior of *g*(*θ*) is the same as in the scenario *θ*_*cw*_ *>* 1 described above: it exhibits the four qualitatively different regimes depending on where the *α*-value lies with respect to 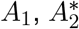 and 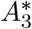. The four regimes are illustrated by the four curves in Fig. 3B. The four regimes are essentially the same as the four regimes in the case *θ*_*cw*_ *>* 1 illustrated in Fig. 3A, but here the range of *θ*-values in which these regimes are realized is restricted to 0 *< θ < θ*_*cw*_. Beyond this range, when *θ > θ*_*cw*_, *g*(*θ*) decreases monotonically as we increase *θ* in all four cases. This is because in this range, the disease-free equilibrium is realized and fungicide treatment does not provide any benefit in terms of disease control, but still incurs a cost. The optimal fungicide coverage, *θ*^∗^ is determined by where *α* lies with respect to 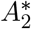. For 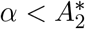, it is best to treat the fraction *θ* = *θ*_*cw*_ of all fields, while for 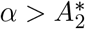 it is best not to treat any fields at all, i.e., to have *θ*^∗^ = 0.

##### B.2.2 Optimal fungicide coverage with resistance

What fraction of fields should be treated with a fungicide to maximize the net return in the presence of fungicide resistance?

###### High-efficacy fungicide and absolute resistance

For simplicity, we first consider the case of a fungicide with the highest efficacy (*ϵ*_*w*_ = 1) and a resistant strain that fully protect the pathogen from the fungicide, i.e., absolute resistance (*ϵ*_*r*_ = 0). As we did above in the scenario without resistance, we investigate the scenario when the system has reached a stable fixed point and analyze the expression for the associated net return given by Eq. (B10) above (Eq. (14) in the main text) But here, 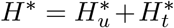 and 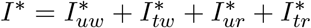. These represent the values of the state variables at a stable fixed point. The system has four fixed points described in Sec. A.3.1 together with their stability conditions. These enter into the expression Eq. (B10) for the net return and determine its dependency on *θ*.

When fungicide resistance confers no fitness cost to the pathogen (*γ* = 0), the fixed point that corresponds to the endemic equilibrium of the resistant strain (FP3 in Sec. A.3.1) is stable (and therefore realized) across the entire range of *θ*-values, i.e. 0 *< θ <* 1. We substitute the expressions for the state variables corresponding to this fixed point into the net return in Eq. (A.3.1) above to find out how *g*(*θ*) depends on *θ* in this regime: *g*(*θ*) = *y* + (1 − *y*)*/* [*R*_0_(1 − *γ*)] − *rθ*, where *R*_0_ = *βN/µ* is the basic reproduction number of the wildtype strain in the absence of fungicide treatment [Eq. (A1)]. In this case, net return, *g*(*θ*), is a decreasing linear function of *θ*, where the slope is given by the fungicide price *f*. This is because in the entire range of *θ*-values, the resistant strain outcompetes the sensitive strain leading to a complete loss of fungicide efficacy. Note that for more expensive fungicides, the net return drops faster as a function of *θ* (Fig. S1A). In this regime, not treating any fields with the fungicide maximizes net return; hence the optimal *θ* is zero.

When fungicide resistance does confer a fitness cost to the pathogen (*γ >* 0), there are three fixed points that can be stable within the permissible range of *θ*-values: (i) the endemic equilibrium of the wild type strain (“wt wins”, FP2 in Sec. A.3.1) at small *θ*-values; (ii) the endemic coexistence of the wild type and the resistant strains (“wt and res coexist”, FP4 in Sec. A.3.1) at intermediate *θ*-values, and (iii) the endemic equilibrium of the resistant strain (“res wins”, FP3 in Sec. A.3.1) at large *θ*-values. The three domains of stability are demarcated by the grey vertical lines in Fig. S1B. The left vertical line represents the transition from FP2 (“wt wins”) to FP4 (“wt and res coexist”) when *θ* is increased and is given by

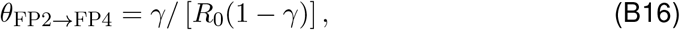

The right vertical line represents the transition from FP4 (“wt and res coexist”) to FP3 (“res wins”) and is given by *θ* = *γ*. In this regime, there can be two qualitatively different scenarios of how the net return depends on *θ*. In the first scenario, the fungicide is sufficiently cheap: it costs less than the yield loss due to disease, *f <* 1 − *y*. In this case, the net return increases over small *θ*-values, reaching a maximum at *θ* = *θ*_*F P* 2→*F P* 4_ and then decreases in a monotonic linear manner as *θ* increases further with the slope given by the relative fungicide price *f* (see in Fig. S1B). In this scenario, the optimal proportion of treated fields that maximizes the net return is given by *θ*_*F P* 2→*F P* 4_ in Eq. (B16) (grey vertical line in Fig. S1B on the left). It is proportional to the fitness cost of resistance *γ* and inversely proportional to the basic reproductive number *R*_0_. Thus, for sufficiently cheap fungicides, there is a non-zero optimal proportion of treated fields, which tends to assume rather small values, as the fitness cost is not expected to exceed 0.3. In the second scenario, the fungicide is more expensive than the yield loss due to disease, *f >* 1 − *y*. In this case, net return decreases monotonically across the entire range of *θ*-values (see Fig. S1B for *f* = 0.5 and *f* = 0.75). This means that the net return is maximum when no fields are treated (*θ* = 0). Thus, in the case of absolute resistance and high-efficacy fungicides, the net return is maximized either at zero or small *θ*-values, and the optimal fraction of treated fields does not depend on the cost ratio *α*. Hence, the economically sound way to deal with fungicide resistance, in this case, is to strongly reduce or stop the fungicide treatment.

###### High-efficacy fungicide and partial resistance

When resistance is partial (0 *< ϵ*_*r*_ *<* 1), the situation is quite different. Here, the qualitative pattern of how *g*(*θ*) depends on *θ* is determined by the cost ratio *α* = *f/*(1 − *y*) [Eq. (15)]. We first focus on a simpler case of no fitness cost (*γ* = 0), and then consider the case when a fitness cost is present (*γ >* 0). How the net return depends on *θ* here is qualitatively similar to the case without resistance discussed above. Since here the resistance is only partial, the fungicide can still suppress the resistant pathogen strain to an extent given by the efficacy parameter *ϵ*_*r*_. If *ϵ*_*r*_ is high enough, then treating a sufficiently high proportion of fields can drive the resistant strain to extinction. This critical proportion of treated fields is given by [Eq. (10) in the main text]

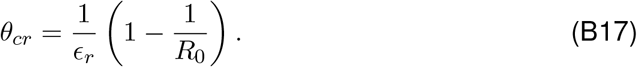

**Figure S1:**
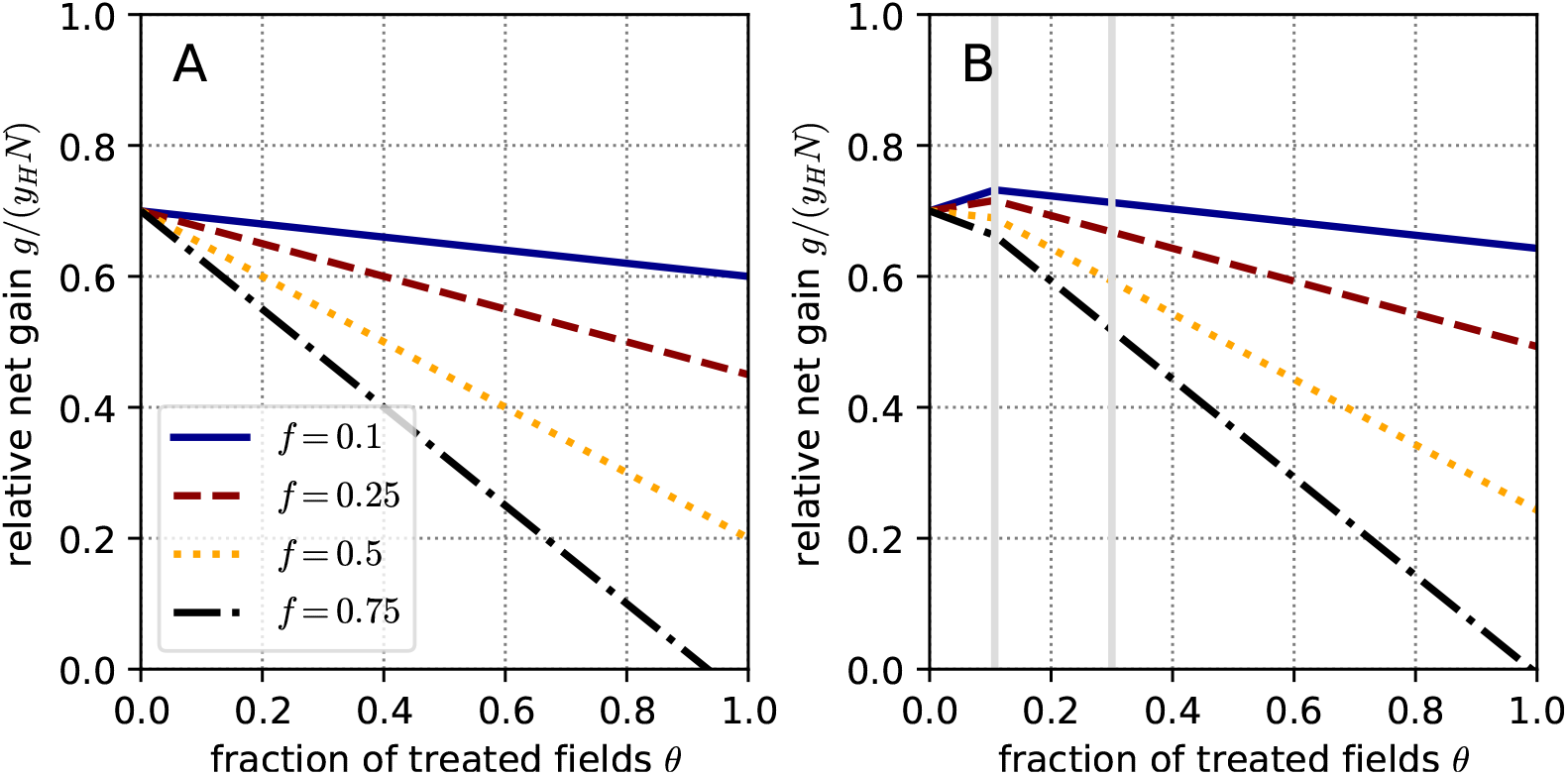
What is the optimal fraction of treated fields in the presence of fungicide resistance (absolute resistance)? Net return *g*(*θ*) is plotted versus the fraction of treated fields *θ* according to Eq. (B10), assuming a high-efficacy fungicide (*ϵ*_*w*_ = 1) and absolute resistance (*ϵ*_*r*_ = 0). Panel A: resistance incurs no fitness cost (*γ* = 0). Panel B: resistance does incur a fitness cost (*γ* = 0.3). The three curves in each panel represent different values of the relative fungicide price *f* : 0.1 (solid blue), 0.25 (dashed red), 0.5 (dotted orange), 0.75 (dash-dotted black). Other parameter values are the same for all curves in both panels: *β* = 0.02, *R*_0_ = *βN/µ* = 4; *N* = 1000, *µ* = 5, *y* = 0.6.

The expression in Eq. (B17) is similar to the critical proportion *θ*_*c*_ given by Eq. (5), but here the entire expression is scaled by the fungicide efficacy against the resistant strain, *ϵ*_*r*_. This difference reflects the fact that the critical proportion here is computed when controlling the resistant rather than the wildtype strain. The associated critical efficacy against the resistant strain above which *θ*_*cr*_ *<* 1 is given by [Eq. (11) in the main text]

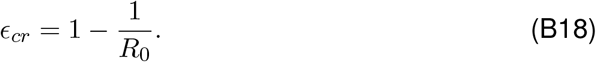

The regime when *θ*_*cr*_ *>* 1 is realized when the fungicide efficacy against the resistant strain is below the critical value, *ϵ*_*r*_ *< ϵ*_*cr*_. In this regime, the only fixed point that is stable and realized across the entire permissible range of *θ*-values is the endemic equilibrium of the resistant strain (FP5 “res wins” in Sec. A.3.2). The other regime, when *θ*_*cr*_ *<* 1 is realized when the fungicide efficacy against the resistant strain is above the critical value, *ϵ*_*r*_ *> ϵ*_*cr*_. In this regime, two fixed points can be stable across the permissible range of *θ*-values: the endemic equilibrium of the resistant strain (FP5 “res wins” in Sec. A.3.2) at low *θ*-values and the disease-free equilibrium (FP1 “disease-free” in Sec. A.3.2) at high *θ*-values. The value of *θ* at which the domain of stability changes from FP5 to FP1 as we increase *θ* is given by *θ*_*cr*_ in Eq. (B17) above.

In each of these regimes, we computed the equilibrium values of the state variables that correspond to stable fixed points and substitute them into the expression for the net return Eq. (B10) to investigate its dependency on the proportion of treated fields *θ*. This dependency is illustrated in Fig. 4, where we distinguished the regimes when *θ*_*cr*_ *>* 1 (panel A) and *θ*_*cr*_ *<* 1 (panel B). The four curves in Fig. 4A and 4B differ only in *α*-values (*α* = *r/*(1 − *y*) Eq. (15)) and represent four qualitatively different scenarios.

**Figure S2:**
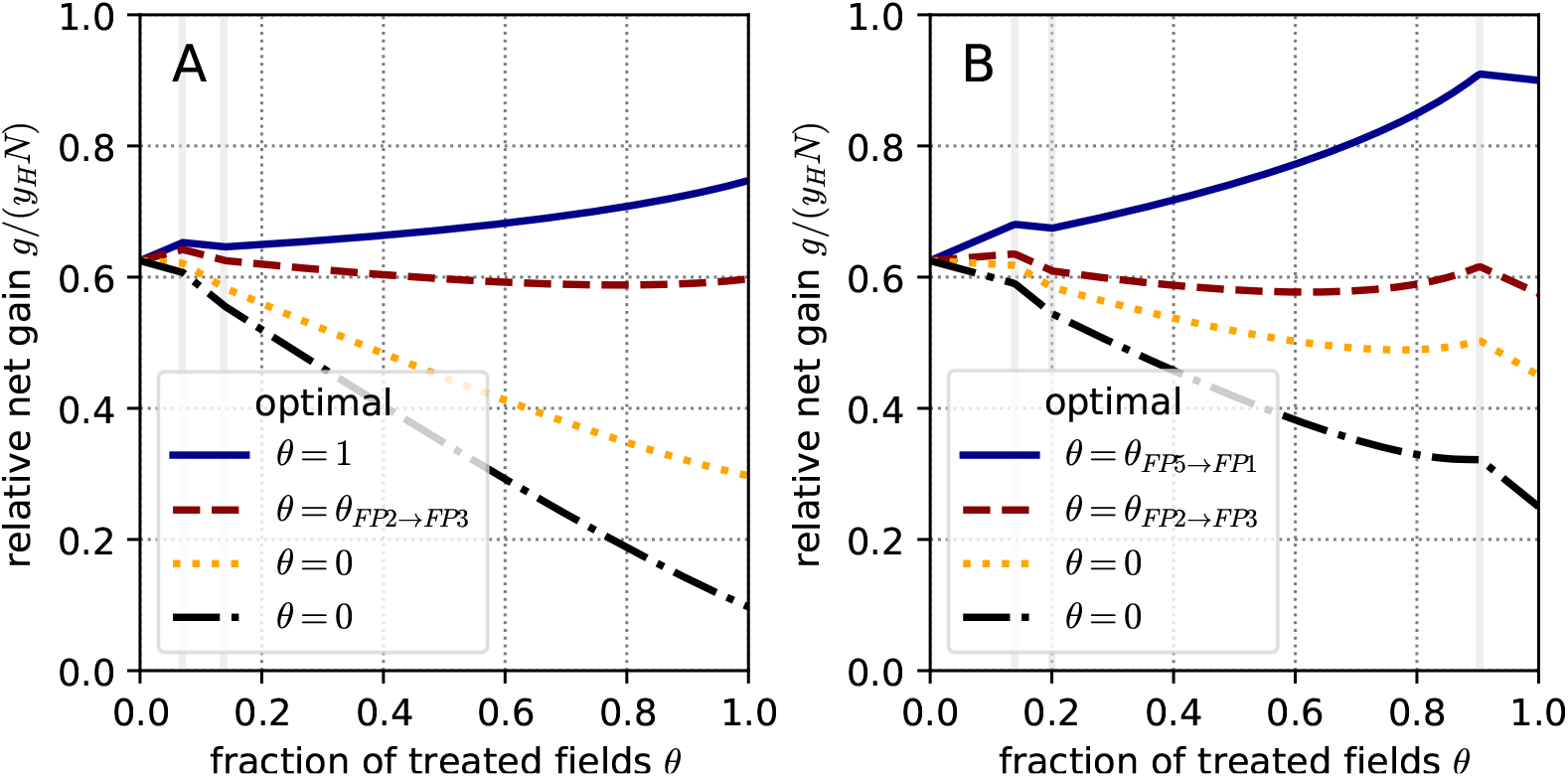
What is the optimal fraction of treated fields in the presence of fungicide resistance (partial resistance with a fitness cost)? Net return *g*(*θ*) is plotted versus the fraction of treated fields *θ* according to Eq. (B10), considering a fitness cost of resistance (*γ* = 0.1), a high-efficacy fungicide (*ϵ*_*w*_ = 1) and partial resistance (0 *< ϵ*_*r*_ *<* 1). Panel A: a low degree of partial resistance *ϵ*_*r*_ = 0.6 *< ϵ*_*cr*_ = 0.75, with values of *α*: 0.1 (solid blue), 0.5 (dashed red), 1.1 (dotted orange), 1.5 (dash-dotted black). Panel B: a high degree of partial resistance *ϵ*_*r*_ = 0.8 *> ϵ*_*cr*_, with values of *α*: 0.1 (solid blue), 0.85 (dashed red), 1.1 (dotted orange), 1.5 (dash-dotted black) Other parameter values are the same for all curves in both panels: *β* = 0.02, *R*_0_ = *βN/µ* = 4; *N* = 1000, *µ* = 5, *y* = 0.5.

Consider the case when the degree of partial resistance is high (*ϵ*_*r*_ *< ϵ*_*cr*_, i.e. *θ*_*cr*_ *>* 1; illustrated in Fig. 4A). Which of the four regimes are realized depends on how the parameter *α* relates to the three constants *B*_1_, *B*_2_ and *B*_3_ that are independent of *θ*, but do depend on the basic reproduction number *R*_0_ and the fungicide efficacy against the resistant strain *ϵ*_*r*_:

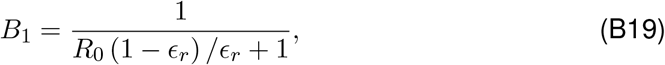

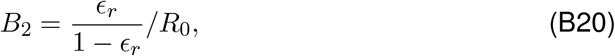

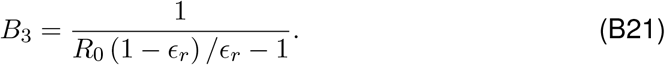

The expressions for *B*_1_, *B*_2_ and *B*_3_ above are the same as the expressions for *A*_1_, *A*_2_ and *A*_3_ in Eq. (B11)-(B13) above, but with *ϵ*_*r*_ instead of *ϵ*_*w*_.

Consider the scenario when the degree of partial resistance is low (*ϵ*_*r*_ *> ϵ*_*cr*_, i.e. *θ*_*cr*_ *>* 1; illustrated in Fig. 4B). As before, which of the four regimes is realized here depends on how the parameter *α* relates to the three constants 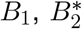 and 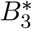 that are independent of *θ*, but do depend on the basic reproductive number *R*_0_ and the fungicide efficacy against the resistant strain *ϵ*_*r*_. The constant *B*_1_ is given by Eq. (B19) and the expressions for 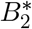 and 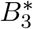 read:

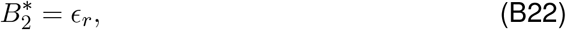

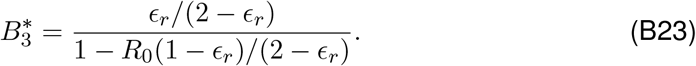

When the fitness cost is present (*γ >* 0), and *θ*_*cr*_ *>* 1, three fixed points can be stable within the permissible range of *θ*-values: the endemic equilibrium of the wildtype strain (FP2 “wt wins” in Sec. A.3.2) at low *θ*-values, the stable co-existence of the wild type and the resistance strains (FP3 “wt and res coexist” in Sec. A.3.2) and the endemic equilibrium of the resistant strain (FP5 “res wins” in Sec. A.3.2) over intermediate *θ*-values. In the different scenario, when *θ*_*cr*_ *<* 1, four fixed points can be stable within the permissible range of *θ*-values: the three fixed points described above plus the disease-free equilibrium (FP1 “disease-free” in Sec. A.3.2) at high *θ*-values.

Based on the stability conditions of FP2, we calculated the *θ*-value at which the domain of stability switches from FP2 to FP3 as we increase *θ*:

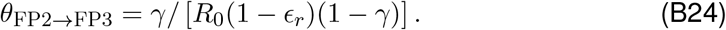

Eq. (B24) represents a generalization of Eq. (B16) for the case of partial resistance. As we did above, we computed the equilibrium values of the state variables that correspond to stable fixed points and substituted them into the expression for the net return Eq. (B10) to investigate its dependency on the proportion of treated fields *θ*. This dependency is illustrated in Fig. S2, where we distinguished the regimes when *θ*_*cr*_ *>* 1 (panel A) and *θ*_*cr*_ *<* 1 (panel B).

The four curves in Fig. S2A differ only in *α*-values (*α* = *f/*(1 − *y*) Eq. (15)) and represent the three qualitative regimes that correspond to different optimal *θ*-values. In the first regime, treating all fields maximizes the net return and hence *θ* = 1 is optimal (see Fig. S2A for *α* = 0.1). This is realized when *α <* 1 and *α < C*_1_ or when *α >* 1 and *α < C*_2_, where

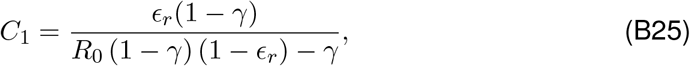

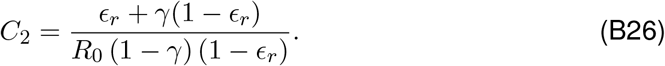

In the second regime, an intermediate value *θ* = *θ*_*F P* 2→*F P* 3_ (given by Eq. (B24), left vertical line in Fig. S2A for for *α* = 0.5) is optimal, which is realized when *α <* 1 and *α > C*_1_ (see Fig. S2A). In the third regime, not treating any fields maximizes the net return and hence *θ* = 0 is optimal (see Fig. S2A for *α* = 1.5 and *α* = 1.1). This is realized when *α >* 1 and *α > C*_2_.

The scenario *θ*_*cr*_ *<* 1 is illustrated in Fig. S2B. Here, there are also three qualitative regimes corresponding to the different optimal values of *θ*. But in this regime, *θ* = 1 is never optimal, because the disease-free equilibrium is stable at high *θ*-values and in this range (to the right from the right-most vertical line in Fig. S2B) the net return given by *g*(*θ*) = 1 − *fθ* decreases linearly with *θ*. The *θ*-value at the point of transition from the domain of stability of FP5 (“res wins”) to FP1 (“disease-free equilibrium”) is given by Eq. (10). Under this scenario, in the first regime, treating the proportion *θ* = *θ*_*cr*_ of fields maximizes the net return. This is realized when *α <* 1 and 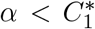 or when *α >* 1 and 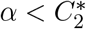, where

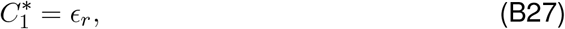

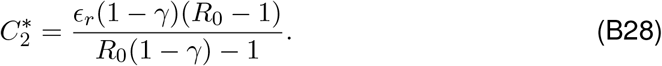

In the second regime, a lower intermediate proportion of treated fields *θ* = *θ*_*F P* 2→*F P* 3_ maximizes net return, which occurs when *α <* 1 and 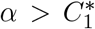. In the third regime, treating no fields maximizes the net return (*θ* = 0 optimal), which is realized when *α >* 1 and 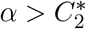.

### C SI. The economic cost of fungicide resistance

#### C.1 Calculation of the economic cost of fungicide resistance

Here, we provide a more detailed mathematical description of the economic cost of resistance, *C*_*R*_. We present analytical expressions for *C*_*R*_ as a function of the relative fungicide price, *f*, in different parameter regimes. This helps us to gain a deeper understanding of the patterns observed in Fig. 6 of the main text.

First, consider a typical dependence of *C*_*R*_ on *f* shown in Fig. 5C in the main text. For low *f* -values, *C*_*R*_ remains constant and is given by

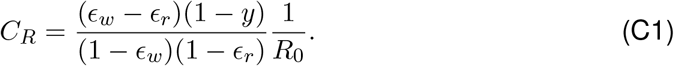

For intermediate *f* -values, *C*_*R*_ decreases linearly with *f* according to

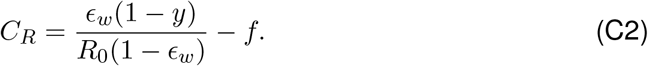

The expressions in Eq. (C1) and Eq. (C2) are valid when the pathogen’s basic reproduction number, *R*_0_, exceeds both of its critical values *R*_0*cw*_ [Eq. (6)] and *R*_0*cr*_ [Eq. (12)]. This means that when all fields are treated, *θ* = 1, the pathogen’s basic reproduction number exceeds one both for the wildtype and the resistant strain.

When the pathogen’s basic reproduction number is high enough (*R*_0_ *> R*_0*cw*_ and *R*_0_ *> R*_0*cr*_) and the fungicide is cheap enough (*f < f*_1_), the economic cost of fungicide resistance is inversely proportional to the pathogen’s basic reproduction number (top left part of Fig. 6B). This can be understood on the basis of the expression for the net return in Eq. (14). The reduction in the net return due to disease is proportional to the fraction of diseased fields *I*^∗^*/N*. This fraction is given by 1 − 1*/* [*R*_0_(1 − *ϵ*_*w*_)] in the absence of resistance, when the endemic equilibrium is reached for the wildtype strain. When the resistance is present and reaches an endemic equilibrium, the fraction of diseased fields is 1 − 1*/* [*R*_0_(1 − *ϵ*_*r*_)]. In order to compute the economic cost of fungicide resistance, we need to subtract the optimal net return with resistance from the optimal net return without resistance. For small *f* -values, when treating all fields maximizes the net return both with and without resistance, the second term in the expression for the net return [Eq. (14)] is the only term that depends on the presence of resistance. Hence, the economic cost of resistance is proportional to the difference between the fraction of diseased fields without resistance and the fraction of diseased fields with resistance. This difference is inversely proportional to the basic reproduction number. The fraction of diseased fields is higher for pathogens with a higher basic reproduction number, *R*_0_, both in the absence and in the presence of resistance. But this increase is higher in the presence of resistance than in the absence of resistance. For this reason, resistance incurs an economic cost. However, because of the nature of the inverse proportionality, at higher *R*_0_-values the difference between the proportion of diseased fields with resistance and without resistance diminishes, and hence the economic cost of resistance is reduced.

When the basic reproduction number is in the range *R*_0*cr*_ *< R*_0_ *< R*_0*cw*_, the wildtype strain can be driven to extinction at an intermediate treatment coverage *θ* = *θ*_*cw*_ *<* 1 (where *θ*_*cw*_ is given by Eq. (5)), but not the resistant strain. For this reason, at small *f* -values the net return without resistance is maximized at this intermediate treatment coverage *θ* = *θ*_*cw*_, while in the presence of resistance treating all fields (*θ*^∗^ = 1) remains the optimum. In this regime, the economic cost of fungicide resistance is given by

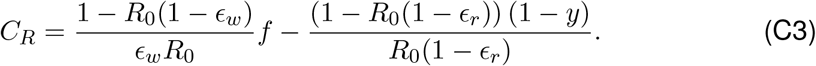

Here, *C*_*R*_ increases linearly as a function of the fungicide price, *f*. This explains the pattern in Fig. 6B we observe for small *f* -values and intermediate *R*_0_-values.

#### C.2 Dependence of the economic cost of resistance on fungicide price and basic reproduction number

In Sec. The economic cost of fungicide resistance of the main text, we focused somewhat more on the scenario when the pathogen’s basic reproduction number exceeds both of its critical values (*R*_0_ *> R*_0*cw*_ in Eq. (6) and *R*_0_ *> R*_0*cr*_ in Eq. (12)), because this scenario appears to be more relevant to fungal crop pathogens. Here, we describe in more detail two alternative scenarios: (i) when *R*_0_ exceeds its critical value for the resistant strain, but not for the wildtype strain (i.e., *R*_0*cr*_ ≤ *R*_0_ *< R*_0*cw*_, and (ii) when *R*_0_ does not exceed its critical value either for the wildtype pathogen strain or for the resistant pathogen strain (*R*_0_ *< R*_0*cw*_ and *R*_0_ *< R*_0*cr*_).

In both scenarios (i) and (ii), *C*_*R*_ increases, reaches a maximum and then decreases reaching zero as a function of *f* (region of the plot corresponding to 1 *< R*_0_ *<* 5 in Fig. 6B), but this pattern arises for somewhat different reasons in each scenario. In scenario (i), the maximum net return without resistance is reached at an intermediate, critical fungicide coverage *θ*^∗^ = *θ*_*cw*_ = (1 − 1*/R*_0_) */ϵ*_*w*_ (Eq. (5)), while the maximum net return in the presence of resistance is reached when treating all fields *θ*^∗^ = 1. As a result, when the fungicide price is increased, the net return without resistance drops at a higher rate compared to the net return in the presence of resistance. This leads to a linear increase in the economic cost of resistance versus fungicide price *f*. But when the fungicide becomes more expensive, the optimal fungicide coverage switches to zero in the presence of resistance. As a result, the net return with resistance remains constant versus *f*, while the net return without resistance continues to decline as we increase *f*. In this range of intermediate fungicide prices, the economic cost of resistance declines linearly with *f* until it reaches zero. Then, the range of high fungicide prices is reached, where no fungicide treatment is economically viable (*θ*^∗^ = 0 both with and without resistance) and hence the economic cost of resistance remains at zero (lower right quadrant of Fig. 6B).

In scenario (ii), in the range of cheap fungicides, optimal fungicide coverage, *θ*^∗^ has an intermediate value both in the absence and presence of resistance. The *θ*^∗^-value is higher with resistance than without resistance because a higher number of diseased fields due to resistance justifies a more intense fungicide treatment. For this reason, the net return with resistance declines faster versus the fungicide price than the net return without resistance. Consequently, *C*_*R*_ increases linearly at low *f* -values. As we further increase *f, C*_*R*_ reaches a maximum, decreases linearly until it reaches zero and remains there as a function of *f* (for the same reasons as in the scenarios described above).

